# Structural Basis for Target Discrimination and Activation by Cas13d

**DOI:** 10.1101/2025.09.12.675955

**Authors:** Chia-Wei Chou, Selma Sinan, Hung-Che Kuo, Carlos Arguello, Daphne Sahaya, Rick Russell, Ilya J. Finkelstein

**Affiliations:** Department of Molecular Biosciences, The University of Texas at Austin, Austin, TX, 78712, USA; Center for Systems and Synthetic Biology, The University of Texas at Austin, Austin, TX 78712, USA

**Keywords:** HEPN, cryo-EM, Cas13, CRISPR RNA, nuclease, mismatch

## Abstract

CRISPR-Cas13d is increasingly used for RNA knockdowns due to its programmability, but off-target RNA binding and cleavage of near-cognate RNAs hinder its broader adoption. Here, we explore the mechanisms of nuclease activation by solving seven ternary cryo-electron microscopy structures of wild-type Cas13d in complex with matched and mismatched targets. These structures reveal a series of active, intermediate, and inactive states that illustrate a detailed activation mechanism. The crRNA undergoes dramatic conformational changes upon target RNA binding, with the helical-1 domain transitioning from an initially docked state with the N-terminal domain to an allosterically switched conformation that stabilizes the RNA duplex. Quantitative kinetics reveal that a single proximal mismatch preserves nanomolar binding affinity but completely abolishes nuclease activity by trapping Cas13d in an inactive state. We identify an active site loop in the HEPN domains that regulates substrate accessibility, with alanine scanning mutagenesis revealing both hypo- and hyperactivated variants. These findings establish the structural basis for Cas13d’s exquisite mismatch surveillance and provide a mechanistic framework for engineering RNA-targeting specificity and activity across HEPN nuclease family members.

## Introduction

Among all class 2 CRISPR-Cas nucleases, Cas13-family enzymes are unique in exclusively targeting and cleaving RNA[1–7]. Cas13 must first search for a target RNA that is complementary to the spacer of the CRISPR RNA (crRNA) via a poorly understood mechanism. After binding the target RNA, the ribonucleoprotein (RNP) is activated to cleave the target (*in cis*), and other RNAs nonspecifically (*in trans*)[1, 5, 8–10]. The ability to exclusively bind and target RNA has spurred intensive development of Cas13 for biotechnology applications, including RNA knockdown, tracking, editing, and nucleic acid detection[3, 11–16]. Despite the broad adoption of Cas13 enzymes across many applications, how Cas13 discriminates between partially matched targets, and how target binding activates the nuclease domain remains poorly understood.

Cas13-family enzymes differ in how they recognize the target RNA, but all use two higher eukaryotes and prokaryotes nucleotide-binding (HEPN) domains to cleave the phosphate backbone [2,5,8,14,17–19]. For example, LshCas13a and BzCas13b, require one or two specific nucleotides, termed the protospacer flanking sequence (PFS), immediately adjacent to the spacer-complement on the target RNA for binding and nuclease activation[1, 3, 4, 14, 15, 18, 20, 21]. The PFS may help these enzymes to identify the matched target RNA, akin to the protospacer adjacent sequence (PAM) for DNA-cleaving CRISPR enzymes[22, 23]. By contrast, Cas13d-family enzymes have a surprisingly high specificity for their target RNAs despite not requiring any PFS for target recognition[3, 12, 24, 25]. Moreover, Cas13d has a high binding affinity to partially matched targets *in vitro*, but these sequences only weakly activate the HEPN domains[26, 27]. Thus, Cas13d is frequently used for RNA knockdown, editing, and diagnostics applications[28–34].

Here, we investigated the molecular mechanisms of Cas13d mismatch surveillance and nuclease activation using a combination of cryo-EM structural analysis and quantitative kinetics. We describe seven Cas13d ternary complex structures, five of which we solve with partially matched targets. These structures revealed two novel states: an intermediate state that precedes the active complex, and an inactive state that occurs when activation fails. A mismatch proximal to the direct repeat constrains the Cas13d complex in an inactive conformation. An active site loop (ASL) regulates Cas13d nuclease activity by interfering with the substrate-binding pocket. Alanine scanning mutagenesis across the ASL identified key residues that further tune nuclease activity. Taken together, our structural and quantitative kinetics data unveil a new activation pathway for Cas13d. More broadly, this study establishes the structural and mechanistic foundation for improved off-target prediction and rational engineering of Cas13d variants with tunable RNA binding and cleavage activity.

## Results

### Cas13d binding and cleavage are uncoupled at mismatched targets

We previously showed that Eubacterium siraeum DSM 15702 (Es)Cas13d nuclease activity is sensitive to mismatches positioned at critical sites along the target RNA[16, 26]. Such mismatches inhibit cleavage without significantly impairing target RNA binding. To better understand the mechanism of this mismatch-dependent nuclease inactivation, we first tested RNA binding and cleavage using quantitative biophysical approaches and a matched target (MT) RNA (**Fig. S1**). Cleavage of a P^32^-labeled RNA substrate showed a strong preference for UU, with a secondary preference for AU dinucleotides (**Fig. S2A**). Therefore, all subsequent experiments use a 10-nt substrate with two central uridines. Substituting both uridines with deoxyuridines (dU) inhibited nuclease activity. Substituting U_5_ with dU did not inhibit nuclease activity. However, a dU_4_ inhibited all cleavage activity, indicating that this 2’-OH participates in the reaction mechanism (**Fig. S2B**).

Next, we investigated Cas13d nucleolytic activity when the RNP binds a partially matched target RNA (**Fig. 1A**). Consistent with our earlier study, a C→A substitution in the 4^th^ position (C4A) of the target RNA largely inhibited RNA cleavage (**Fig.1B**,**C**). By contrast, a U10G mismatch resulted in moderate cleavage. Importantly, both mismatched target RNAs have nearly identical equilibrium binding constants that are dominated by the on-rates, *k*_*on*_ (**Fig. 1C**)[1, 2, 5, 35, 36]. Taken together, these data indicate that partially matched target RNAs bind but fail to activate the Cas13d nuclease.

**Figure 1:**
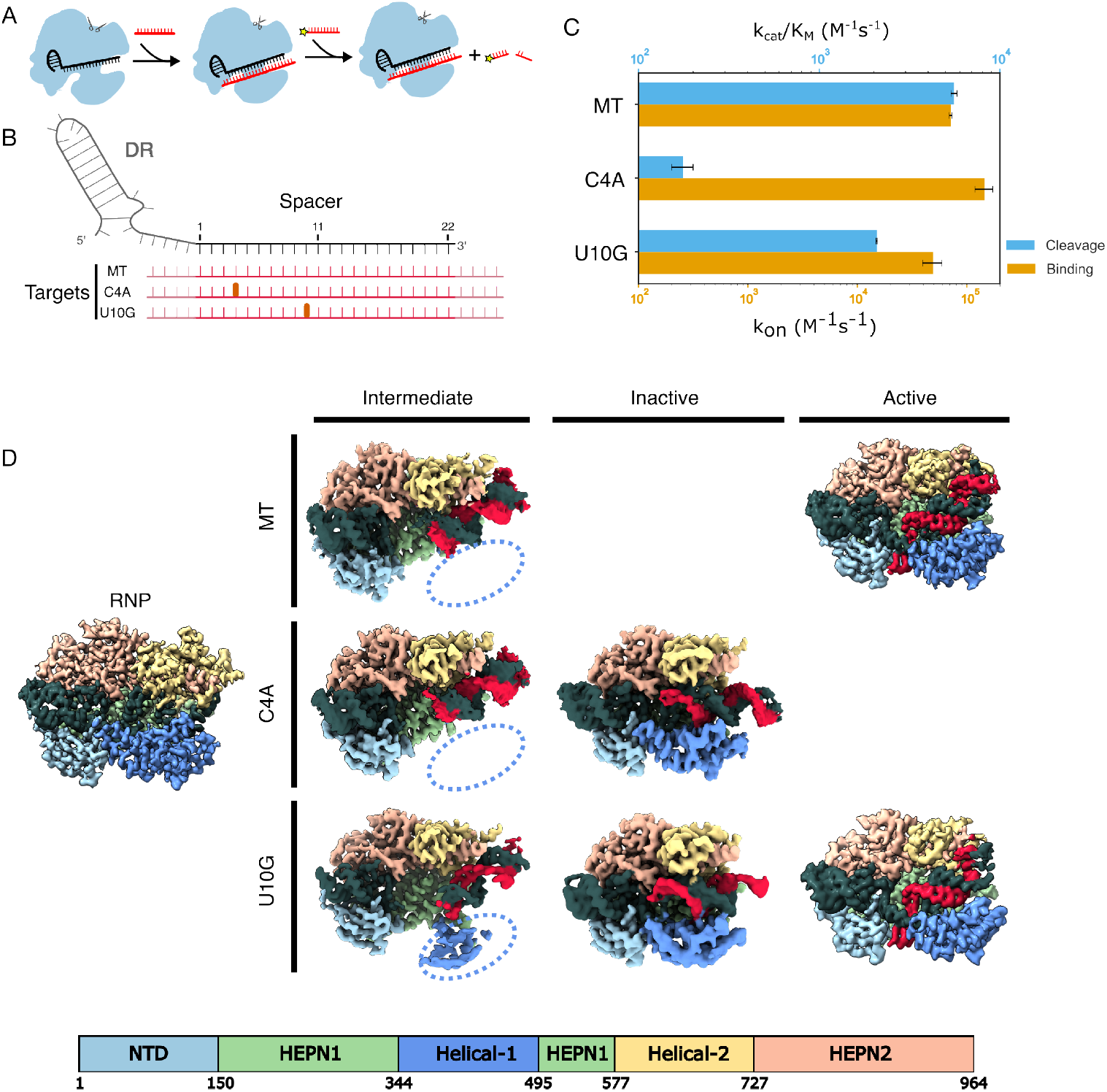
Mismatched RNA targets abrogate RNA cleavage and induce new Cas13d states. (A) Schematic of Cas13d cleavage assay. Cleavage of a P^32^-labeled RNA substrate is triggered by the binding of an RNA target complementary to the crRNA. Black: crRNA, red: target RNA in red, yellow: P^32^. (B) Diagram of the mismatched targets investigated in this study. Red: target RNA, thick ticks: mismatches at the 4th and 10th positions, pink: 5’ and 3’ flanking sequences. (C) Second-order cleavage rates (blue bars) of a *trans* substrate when Cas13d is activated by a matched target (MT), or mismatched targets. Orange bars: on-rates (*k*_*on*_) measured for the same substrates. Error bars: mean ± S.D. from at least three biological replicates. (D) Cryo-EM density maps of seven Cas13d tertiary structures with the indicated mismatched target RNAs. Dashed oval: missing Helical-1 domain density.

### Two new Cas13d ternary complexes revealed by partially matched target RNAs

To elucidate the molecular basis for Cas13d nuclease activation, we determined cryo-EM structures of a binary WT Cas13d complex, as well as ternary complexes with a matched target, a C4A mismatch, or a U10G mismatch (**Fig. 1D** and **Figs. S3-S6**). The binary structure closely resembled that of a previously-published Cas13d (PDB:6E9E) with C*α*-C*α*-RMSD of 1.33 Å (**Fig. S7**)[18]. A central hallmark of this structure is a kinked crRNA and a Helical-1 domain that is docked and interacting with the N-terminal domain (NTD). The 5’-end of the crRNA spacer is clamped between the Helical-1 and Helical-2 domains, but the 3’-distal end of the spacer is solvent-exposed and wedged between the Helical-2 and HEPN1 domains. These results are broadly consistent with earlier studies and support a model where the target RNA first hybridizes with the 3’-distal end of the crRNA spacer[26].

The ternary structures also revealed a new conformation with a kinked crRNA-target RNA duplex and a flexible Helical-1 domain (**Fig. 1D, intermediate state**). Notably, the target RNA is not resolved after the 23rd nucleotide in our models, indicating that it remains flexible. Three lines of evidence suggest that this complex represents an intermediate “checkpoint” that precedes HEPN nuclease domain activation and subsequent cleavage of the target RNA. First, this state is populated in about half of the matched target complexes, as well as half of the C4A and one-third of U10G complexes in the cryoEM single-particle analysis, indicating that this is an intermediate along the nuclease activation pathway(**Figs. S3-S5**). Second, the crRNA in this state extends towards the 3’ end to be more exposed to the solvent and form a complete RNA duplex. Although the crRNA is slightly kinked, as in the matched target Cas13d structure, the RNA duplex inclines toward the crevice between the Helical-2 and HEPN1 domains instead of binding between the Helical-2 domain and the HEPN2 domain (**Fig. 2**). Finally, the U10G structure captures some density for the flexible Helical-1 domain, indicating that it is no longer docked with the NTD (**Fig. 1D, dashed oval**). Rearranging NTD-Helical-1 interactions is essential for establishing the active ternary structure (see below). However, the active site is conformationally similar to the binary complex, indicating that this state is not yet poised for target RNA cleavage. Thus, we conclude that this state is a transient but stable intermediate in the Cas13d activation pathway.

**Figure 2:**
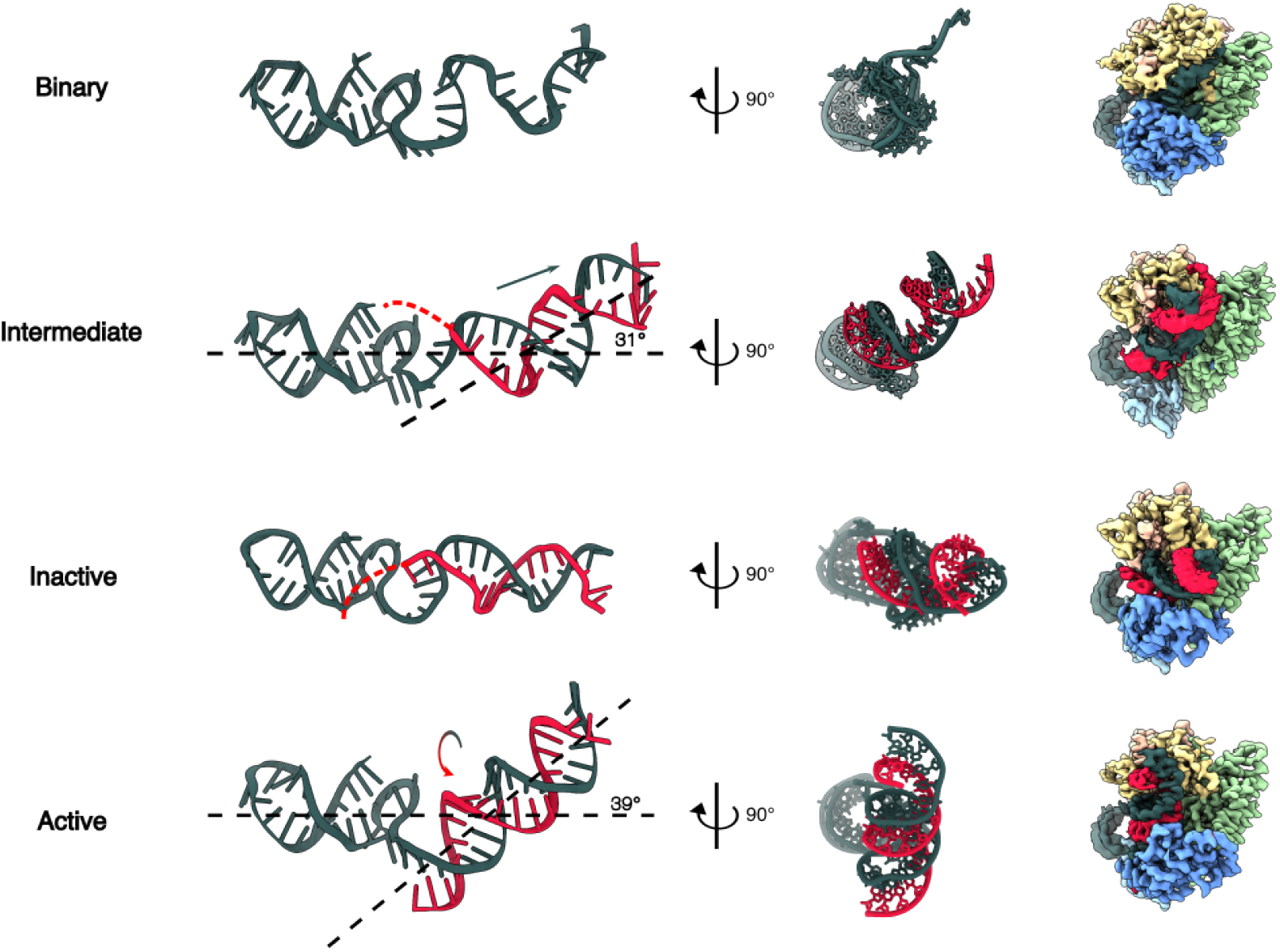
Conformational changes in RNA duplexes across different states. Left: Front view of crRNA and RNA duplexes across all states. Middle: Side view of crRNA and RNA duplexes across all states. Right: Side view of cryo-EM density maps across all states.

The C4A and U10G RNPs also reveal a second conformation, which we term an “inactive” state based on our quantitative kinetic assays and structural considerations (**Fig. 1D, middle row**). This conformation is characterized by a linear crRNA-target duplex with a Helical-1 domain stably anchored to the NTD. The RNA duplex stops at position 5 in the C4A complex, indicating that this mismatch locks the complex into a state that cannot mobilize Helical-1 for subsequent conformational changes. We defined the linear conformation as an inactive state of Cas13d for the following reasons. First, we only observed this state with mismatched ternary complexes that ablate or drastically reduce nuclease activity (**Fig. 1C**). Second, this state has a straight helix in the HEPN2 domain, in contrast to the kinked conformation observed in the active state (**Fig.S8**)[18]. Third, the active site of Cas13d has a conformation similar to that of binary Cas13d. Specifically, the Helical-1 domain still interacts with the NTD, similar to the binary complex (see below).

We also solved the structure of WT Cas13d in complex with a matched target RNA. The overall structure is very similar to the previously published catalytically-inactive Cas13d (R295A, H300A, R849A, H854A), termed dCas13d (**Fig. S7**)[18]. The overall C*α*-C*α*-RMSD between our structure and the ternary dCas13d structure (6E9F) is 0.7Å. However, steric clashes between the native residues in the HEPN1 and HEPN2 domains reduce the local alignment to 2.6Å in the nuclease active site and will be discussed below. As noted above, one of the two U10G structures is identical to that of the matched target, with one exception. To accommodate the U10G mismatch, the crRNA and target RNAs form a non-A-form RNA duplex at the distal part of the crRNA and a canonical A-form duplex in the proximal crRNA region. This makes the N-terminal HEPN1 *α*-helix bent relative to the active MT structure. Below, we discuss the structural implications of these distortions on HEPN domain activation.

### Global domain motions required for target RNA recognition and nuclease activation

The crRNA must undergo global rearrangements as Cas13d transitions from the binary to the active state (**Fig. 2 and Fig. S9**). In the binary state, the crRNA is bulged towards its 3’-end and makes the distal spacer more exposed to the solvent. In the intermediate state, the crRNA engages with the target RNA and extends toward the 3’ end. The Helical-1 domain undocks from the NTD to accommodate further pairing of the spacer and target RNAs. The nascent RNA duplex stays in the cleft that was previously occupied by the spacer in the binary state. RNA base pairing stops at position 5 in the intermediate state, indicating that Cas13d imposes an energy barrier to further base pairing in the proximal positions. The weak EM density around the proximal spacer suggests that the RNA is heterogeneous or dynamic in this region. In the inactive state, the RNA duplex does not continue into the proximal bases, allowing the Helical-1 domain to stabilize the partial RNA duplex. This tilts the RNA duplex ~30°relative to the plane. In the matched target structure, RNA base pairing extends along the entire spacer. This forces the crRNA to rotate clockwise and insert between the NTD and Helical-1 domains. The RNA duplex tilts back ~40°and moves forward from the Helical-2 /HEPN1 cleft to the HEPN1 /HEPN2 crevice, opening the original charge surface on the HEPN1 and Helical-2 domain for substrate binding (**Fig. S10A**). Mutations of charged residues in the HEPN1 domain reduced cleavage activity by 30% to 70%, indicating that this surface is critical for substrate binding. (**Fig. S10B, C**)

### The crRNA undergoes a dramatic conformational change upon nuclease activation

Extensive interactions between the Helical-1 and NTD domains in the inactive state sterically block the formation of the RNA duplex (**Fig. 3A**). Helical-1 residues R435 and D428 form hydrophilic interactions with G83 and R84 in the NTD. However, the active state disrupts these interactions to make room for the RNA duplex, which forms hydrogen bonds with R84, N86, N377, and R435 (**Fig. 3B**). Arginines R435 and R84 interact directly with target RNA bases and are essential for cleavage. N86 interacts with A (−2) in the direct repeat and N532 in the HEPN1 domain in the inactive state while it interacts with the target backbone in the active state. N377 has no interaction in the inactive state but binds in the minor groove of the proximal RNA duplex. Disrupting any of these four residues via alanine substitutions significantly reduces cleavage but not the equilibrium binding constant (**Fig. 3C**). In sum, proximal mismatches disrupt nuclease activation by preventing the RNA duplex from reorganizing NTD-Helical-1 interactions.

**Figure 3:**
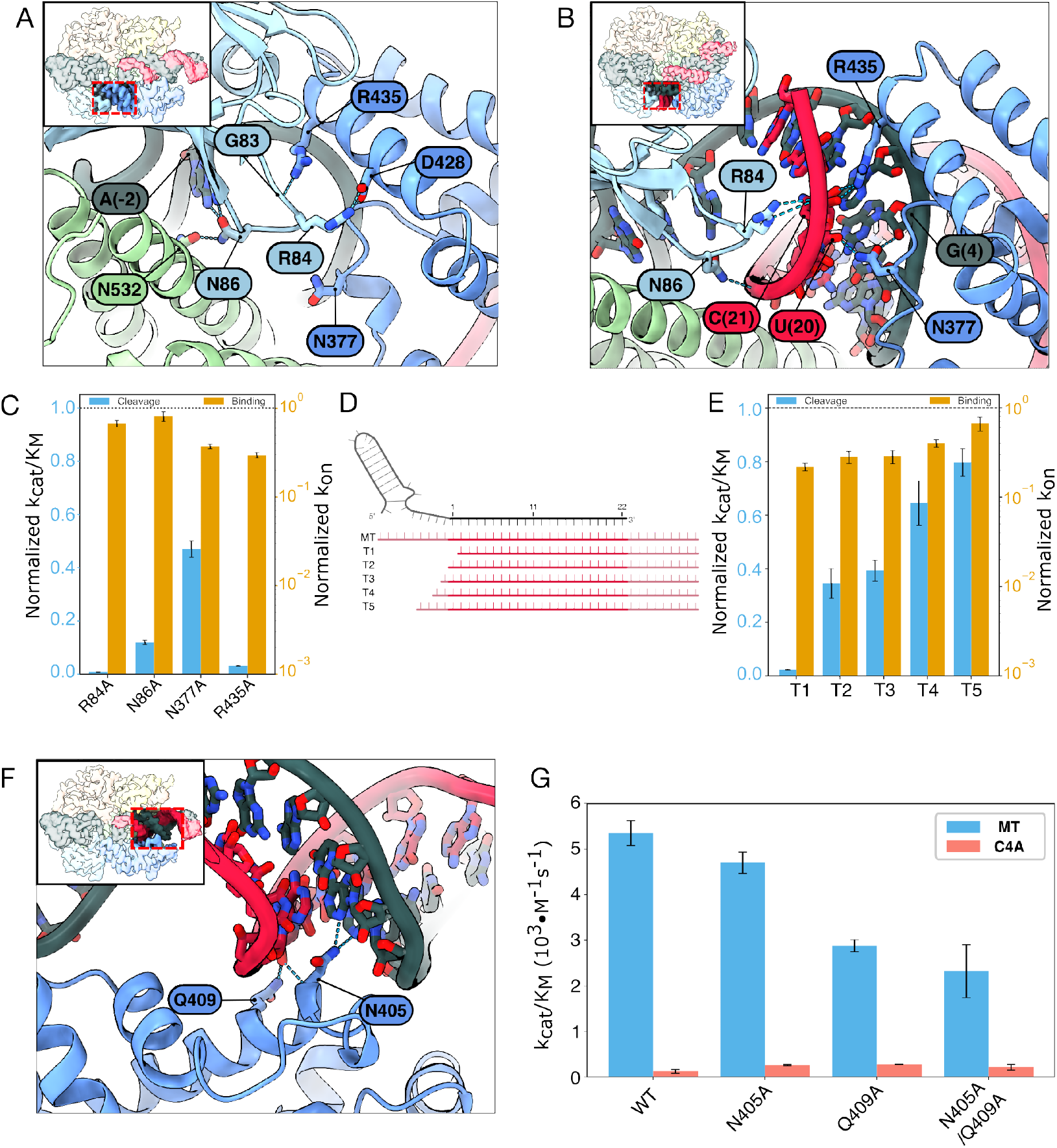
RNA-Cas13d interactions that change between the inactive and active states. Enlarged view of NTD-Helical-1 domain interactions in the (A) inactive and (B) nuclease-active states. Key changes between the two states include disruption of R435-G83 and D428-R84 interactions, formation of new RNA-protein contacts with R84, N86, N377, and R435, and displacement of the Helical-1 domain to accommodate the extended RNA duplex. (C) Second-order cleavage rates of a *trans* substrate by Cas13d mutants, normalized to the WT Cas13d rates shown in the dashed line. Error bars: S.D. from three biological replicates. (D) Schematic of a series of progressively truncated RNA targets, labeled T1-T5. Light red: 5’ and 3’ flanking sequences. (E) Secondorder cleavage rates activated with the indicated RNAs. The rates are normalized to the extended MT RNA (dashed line). Error bars: S.D. from three biological replicates. (F) Enlarged view of N405/Q409 interactions with crRNA positions 10 and 11 in the inactive state that stabilize the linear RNA duplex conformation. (G) Second-ord1e7r cleavage rates of N405A and Q409A mutants with MT and C4A targets, showing partial rescue of C4A cleavage activity. Error bars: S.D. from three biological replicates.

Based on these mutagenesis studies, we hypothesized that the target RNA duplex must also be sufficiently long to disrupt these interactions, even when it is fully matched to the crRNA. To test this hypothesis, we measured the cleavage rates and binding affinities for a series of truncated MT RNAs (**Fig. 3D, E**). Remarkably, even a single nucleotide truncation relative to the spacer completely ablates nuclease activity (T1). Truncations T2-T5, which partially extended the target RNA, also showed reduced activation that was proportional to the length of the truncation. These results confirm that the RNA duplex must extend beyond the spacer to fully liberate the Helical-1 domain for nuclease activation. Moreover, nuclease activation can be tuned *in vitro* and in cells via judicious selection of mismatches and target RNAs (see Discussion).

The RNA duplex is linear in the inactive state. This structure is partially stabilized by hydrogen bonds between N405/Q409 and positions 10 and 11 of the crRNA. These interactions do not exist in the active and intermediate MT and U10G structures (**Fig. 3F**). Thus, we tested whether the alanine substitutions N405A and Q409A can re-activate cleavage of the C4A target. Cleavage of the C4A target increased 1.6-fold relative to WT Cas13d for both N405A and Q409A, with no synergistic effect for the N405A/Q409A double mutant (**Fig. 3G**). However, both mutants also reduced the cleavage rate of the MT substrate by up to 40%, revealing a sharp tradeoff between enzyme promiscuity and nuclease activation.

### An active site loop regulates trans-substrate accessibility by the HEPN domains

To understand how the HEPN nuclease is activated upon target RNA binding, we solved all structures with the fully WT active site residues. These structures are in contrast with prior Cas13d studies that used the nuclease-inactive Cas13d (H854/H300A/R295A/R849A) [18, 37]. Remarkably, the binary, intermediate, and inactive states retained the same overall active site architecture (**Fig. 4A, B**). However, the HEPN2 *α*-helix moved 2.9Å from the binary to the active state.

**Figure 4:**
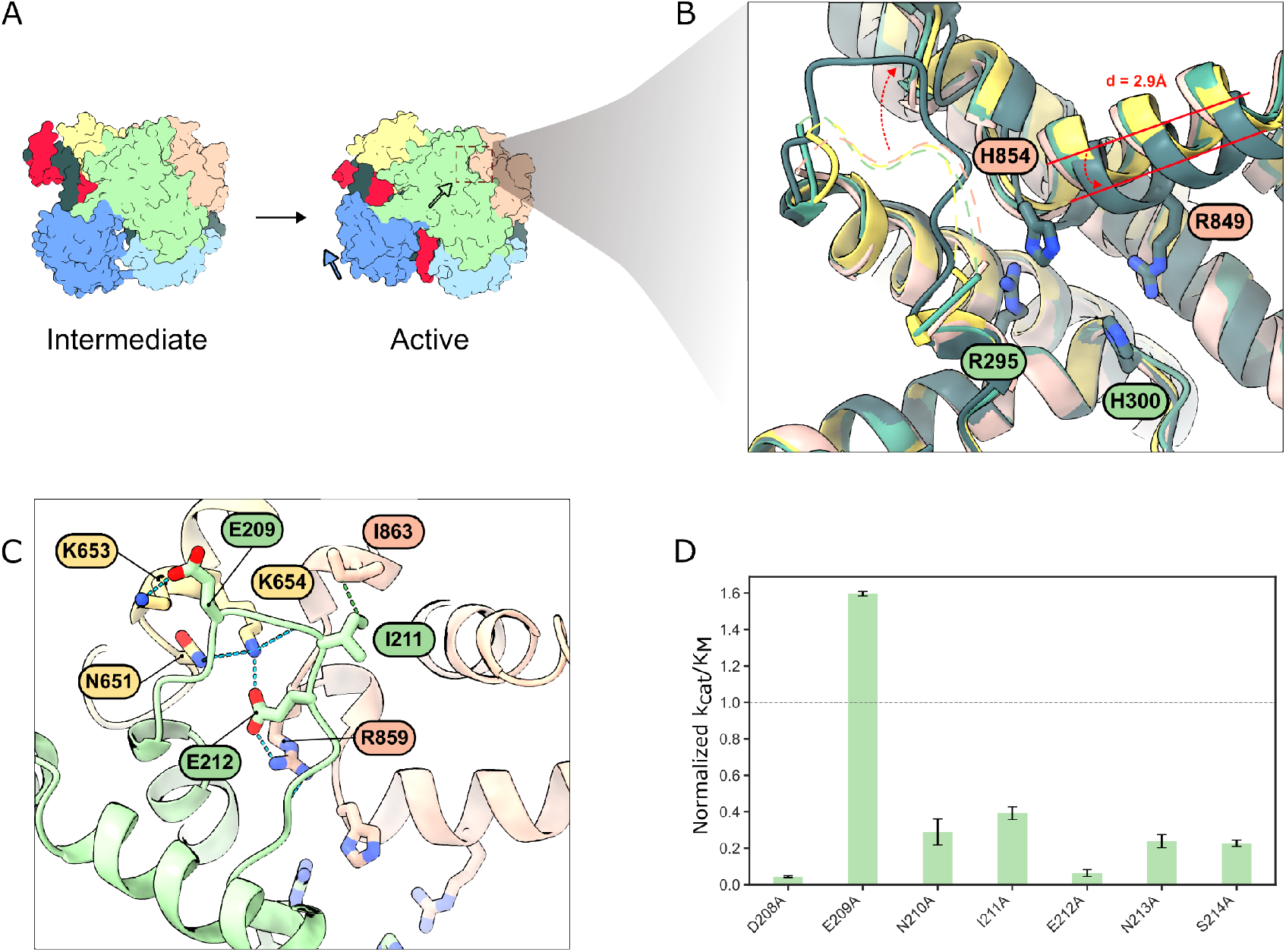
Stabilization of the active site loop regulates *trans* cleavage rates. (A) A back view of global domain movements between the intermediate and active states. The arrows indicate global domain movements from the intermediate state to the active state.(B) An enlarged view shows the superposition of the active sites in the binary (pink), intermediate (yellow), inactive (light green), and active (dark green) states. (C) Detailed interactions with ASL residues. Green: HEPN1 domain, pink: HEPN2, yellow: Helical-2. (D) Second-order cleavage rates with the indicated Cas13d mutants, activated with an MT RNA and normalized to WT Cas13d (dashed line). Error bars: S.D. of three biological replicates.

We also resolved a loop near the active site that mainly interacts with the HEPN2 but also weakly interacts with the Helical-2 domain. Because of its proximity to the active site, we termed it the active site loop (ASL, **Fig. 4C**). The ASL is only stabilized by the HEPN2 and Helical-2 domains in the active state. All other reported structures, including our own, do not resolve this loop. AlphaFold models of four additional Cas13d orthologs confirmed that the ASL is a broadly conserved feature of this family of enzymes (**Fig. S11**)[38].

We reasoned that the ASL may regulate substrate access into an electrostatic binding pocket for nucleolytic cleavage (**Fig. S11B**). Nearly all alanine substitutions along the ASL significantly reduced cleavage activity relative to wild-type enzymes, suggesting that stabilization of the ASL-HEPN2 interface is critical for substrate binding. Interestingly, the E209A mutation— the only residue interacting with the Helical-2 domain—enhanced cleavage activity by approximately 1.6-fold (**Fig. 4D**). We hypothesize that E209A may hyperactivate the enzyme by liberating K653 to interact with the RNA electrostatically. We conclude that the ASL regulates accessibility of the RNA for the HEPN active site and that engineering this region may be a fruitful avenue for further hyperactivating Cas13d.

## Discussion

Here, we report the structural basis for Cas13d nuclease activation and mismatch surveillance (**Fig. 5**). To our knowledge, this is the first structural study of any Cas13 family member that retains all the wild-type residues in the HEPN active site. A growing body of evidence suggests that the target RNA first basepairs with the hairpin-distal side of the crRNA [3, 4, 24, 26, 39]. Mismatches between the target RNA and the distal side of the crRNA impede target recognition and reduce the overall RNA binding affinity[24]. Base pairing proceeds toward the proximal end of the crRNA via an intermediate state that weakens the Helical1-NTD interactions in advance of an extending dsRNA. To activate the nuclease, the NTD and Helical-1 domains must separate and make sufficient room for RNA duplex formation. Mismatches towards the middle of the spacer, i.e., the U10G in this study can be partially bypassed via non-Watson-Crick basepairing, as has been observed for Cas9 and other CRISPR-Cas proteins[40–43]. Mismatches near the proximal end fail to disrupt the NTD-helical-1 interactions, locking the RNP in an inactive state. Displacement of the Helical-1 domain is crucial for allosteric activation of the catalytic site by bringing the two HEPN domains in closer proximity than in the inactive state. This multi-step verification process ultimately activates the nuclease and explains Cas13d’s ability to discriminate against partially complementary off-target RNAs [25, 26].

**Figure 5:**
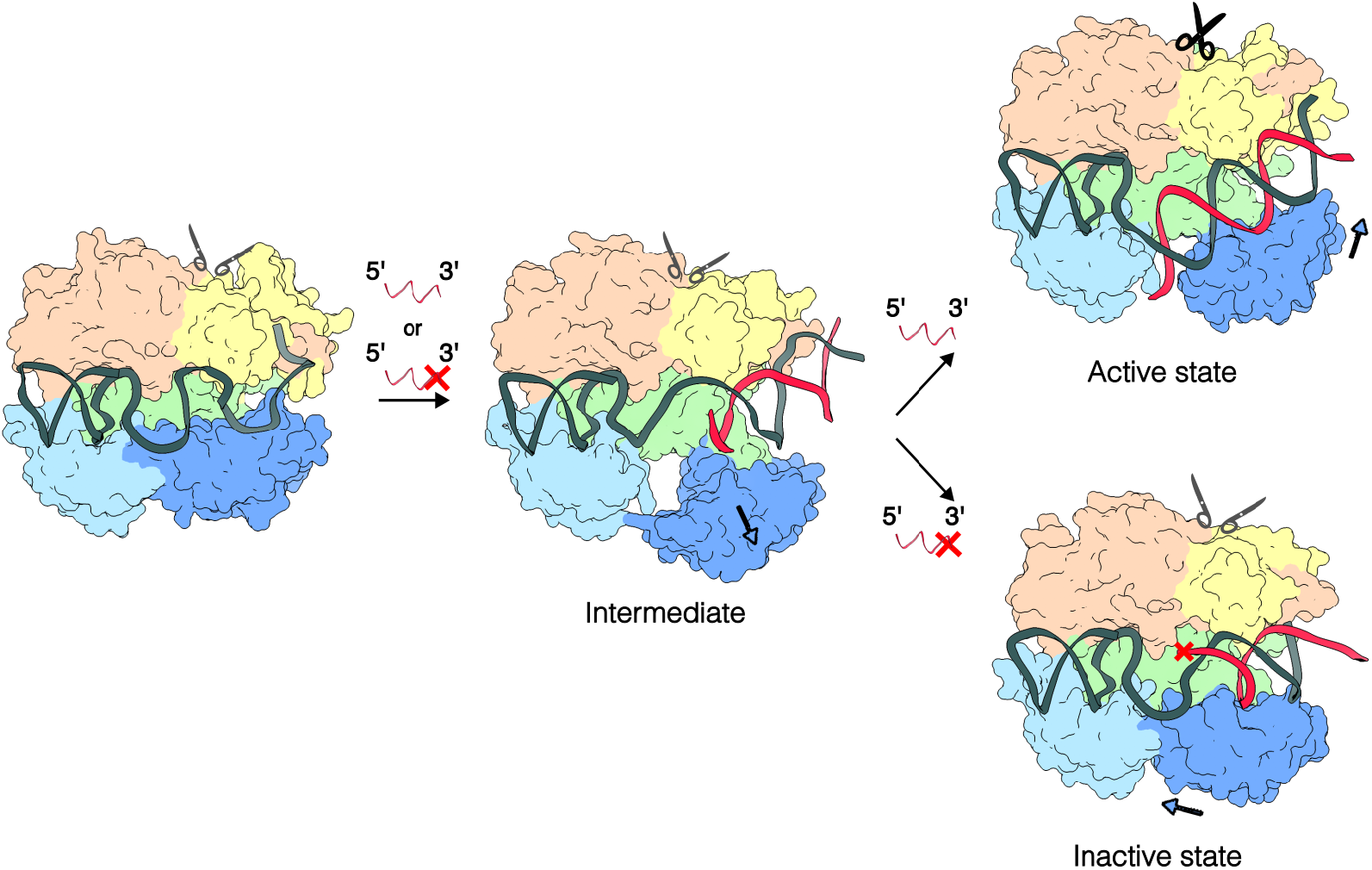
Proposed Cas13d activation model. Cas13d Ribonucleoprotein (RNP) recognizes distal mismatches with low binding affinity, which may prevent the initiation of RNA duplex formation (target with red cross). By contrast, an RNA with a partial match at the distal end can initiate duplex formation, progressing the ternary complex to an intermediate state. The formation of a proximal RNA duplex must separate multiple contacts between the NTD and Helical-1 domains, creating an energy barrier for mismatch surveillance. A perfectly matched target activates Cas13d’s HEPN nuclease domain (top, scissors). Target RNAs with a proximal mismatch trap Cas13d in an inactive state (broken scissors).

Cas13d’s high specificity for its target RNA may have evolved to minimize auto-immunity in bacteria. Unlike other type VI CRISPR nucleases, Cas13d doesn’t require a protospacer flanking sequence (PFS)[4]. This lack of stringent PFS requirements lets Cas13d target almost any RNA. By contrast, Cas13a requires a non-G PFS; BzCas13b favors non-C 5’-PFS and 3’PFS of NNA or NAN[1, 2, 5]. Further, Cas13d does not require an anti-tag, further broadening its target range[24, 44]. Cas13d activation is toxic to cells, leading to growth inhibition and potential cell death[45]. Thus, Cas13d must rely on the overall complementarity between the crRNA guide and the target RNA, with mismatches blocking nuclease activation (**Fig.5**).

HEPN nucleases are metal-ion independent and catalyze RNA cleavage via a 2’-O-transesterification mechanism[5, 46–49]. However, prior reports concluded that Cas13 strictly requires a divalent metal ion [1, 5, 8]. This suggests that either the Cas13 HEPN domain has evolved a catalytic requirement for metal ions or that the metal ions stabilize the substrate for metal-independent cleavage. We also observed active site electron density that is consistent with a Mg^2+^ ion coordinated by two conserved histidines (H300 and H854; **Fig. S12A**). RNA cleavage requires Mg^2+^ for rapid substrate degradation, with no cleavage after 3 hours in the absence of Mg^2+^. While underscoring the importance for Mg^2+^, we cannot distinguish between these two mechanisms (**Fig. S12B**). Future studies will need to focus on the role of metal ions in HEPN nuclease activation.

The inactive, intermediate, and active Cas13d structures only differ by a 2.9Å movement between HEPN-1 and HEPN-2 domains. These results are in contrast to prior structural studies in Cas13bt3, where the HEPN domains shift together by about 24Å[17, 19]. LbuCas13a also shows large rearrangements in the nuclease active site, with HEPN1 moving 6.5–9.6 Å toward the HEPN2 domain after the crRNA-target RNA duplex forms[50]. We speculate that in Cas13d, the ASL regulates access to the HEPN nuclease active site. This mechanism is reminiscent of how some proteases and kinases regulate access to their active sites[51, 52]. Indeed, a single mutation (E209A) hyper-activates the Cas13d nuclease, possibly by opening the positively charged substrate-binding pocket and pulling the HEPN-1 and HEPN-2 domains together. Further engineering of the ASL may boost RNAse activity for both diagnostics and RNA editing applications.

Non-specific nuclease activation and subsequent indiscriminate RNA cleavage has limited Cas13-family nuclease use for biotechnological applications cells[53–55]. Widespread RNA degradation can lead to cellular stress, altered gene expression, and cell death. In RNA diagnostics, this collateral activity can result in false positives or overly sensitive detection, complicating experiment interpretation. To address these issues, researchers have explored strategies to modulate Cas13 activity. One approach is to attenuate nuclease activity via Cas13d engineering[54]. A simpler approach is to introduce programmed mismatches between the crRNA and its target to reduce the number of active nucleases, thereby minimizing non-specific effects[24]. Our study provides a mechanistic basis for installing such mismatches. A proximal mismatch can effectively down-regulate nuclease activity without changing RNA binding affinity. In contrast, for applications requiring high sensitivity such as in viral RNA detection, hyperactive mutations in ASL hyperactivate the nuclease. Future protein engineering studies will focus on further engineering Cas13d-based tools for mammalian systems, particularly in therapeutic contexts where precise but rapid RNA targeting is crucial.

## Materials and Methods

### Protein cloning and purification

Oligonucleotides were purchased from IDT. Wild type EsCas13d was cloned into a pET19 vector with an N-terminal 6*×*His-TwinStrep-SUMO purification tag to generate plasmid pIF1023 (Table 1)[26]. Nuclease-dead dCas13d was cloned by introducing R295A, H300A, R849A and H854A mutations into plasmid pIF1024 via the Q5 Site-Directed Mutagenesis Kit (NEB)(Table 1). Variant Cas13d proteins were cloned using the same kit.

To purify the wild-type protein, pIF1023 was transformed into *E. coli* BL21 (DE3) cells. Cells were induced with 0.5 mM isopropyl *β*-D-1-thiogalactopyranoside (IPTG) at OD_600_~0.6-0.8, and grown at 18 °C for 16-20 hours. Cells were pelleted and stored at −80 °C. For purification, pellets were thawed and resolubilized in Lysis Buffer (50mM HEPES pH 7.4, 500 mM NaCl, 1 mM EDTA, 5% glycerol, 1 mM DTT, EDTA-free protease inhibitor cocktail (cOmplete), RNase-free DNase I (NEB), salt active nuclease (NEB)). The resuspended pellet was sonicated, and cell debris was pelleted at 4 °C via centrifugation at 20,000 g for 45 mins. The supernatant was incubated with Strep-Tactin resin (IBA) and further washed with Wash Buffer (50 mM HEPES pH 7.4, 500 mM NaCl, 1 mM DTT, 5% glycerol) and eluted with elution buffer (50 mM HEPES pH 7.4, 500 mM NaCl, 1 mM DTT, 10% glycerol 7.5 mM D-dethiobiotin (DTT)). Elutions were treated with homemade SUMO protease overnight at 4 °C. ApoCas13d was further purified by size exclusion chromatography using a Superdex 200 Increase (GE healthcare) in SEC Buffer1 (50 mM Tris pH 7.5, 100 mM NaCl, 5% glycerol, 1 mM MgCl_2_ and 1 mM DTT) for structural studies and in SEC Buffer2 (25 mM Tris pH 7.5, 200 mM NaCl, 5% glycerol, 1 mM MgCl_2_ and 1 mM DTT) for kinetic assays. Fractions were further pooled, concentrated, flash frozen, and stored at −80 °C. All variants were purified using the same protocol.

### Binary and ternary complex assembly and purification

Cas13d-RNA complexes were assembled by incubating purified Cas13d with crRNA (IDT) at a 1:3 molar ratio in RNP Formation Buffer (50 mM Tris-HCl pH 7.5, 100 mM NaCl, 6 mM MgCl_2_, 1 mM DTT) at 37 °C for 1 hour. The binary complex was separated from excess crRNA by further purified by size-exclusion chromatography using a Superdex 200 Increase 10/300 GL column (GE Life Sciences) equilibrated with SEC buffer (50 mM Tris-HCl pH 7.5, 150 mM NaCl, 1 mM MgCl_2_, 10% glycerol, 1 mM DTT). RNP-containing fractions were pooled and concentrated with Amicon Ultra-15 centrifugal filters (30 kDa MWCO, Millipore). For ternary complex formation, the binary RNP was incubated with target RNA (IDT) at a 1:5 molar ratio and purified by SEC under identical conditions. The purified complexes were concentrated, flash-frozen in liquid nitrogen as single-use aliquots, and stored at −80 °C.

### Nuclease assays

The Cas13d-crRNA complex was assembled daily for experiments using purified Cas13d and a crRNA purchased from IDT (Table 2). Assembly reactions were carried out by incubating Cas13d with a crRNA concentration that varied and was lower than that of Cas13d for 30 minutes at 37 °C in a buffer containing 50 mM Tris-HCl, 100 mM NaCl, 6 mM MgCl_2_, and 1 mM DTT.

For trans cleavage reactions, the target RNA (10 nt) was 5’-radiolabeled with [*γ*-32P] ATP (PerkinElmer) using T4 polynucleotide kinase (New England Biolabs). The Cas13d-crRNA complex was incubated with varying concentrations of different sizes of target RNA for 30 minutes at 37 °C in a reaction buffer containing 50 mM Tris-HCl, 150 mM NaCl, 6 mM MgCl_2_, 1 mM DTT, and 0.2 mg/ml molecular-grade BSA (Table 2). Trans cleavage reactions were initiated by adding a trace amount of labeled target RNA to varying concentrations of the ternary complex. At different time points, 2 µL samples were quenched in 4 µL of denaturing solution (20 mM EDTA, 90% formamide, 1.5 mg/ml Proteinase K, and 0.05% xylene cyanol). Samples were analyzed by denaturing PAGE (20% acrylamide, 7 M urea). The gels were exposed to a phosphorimager screen overnight, scanned with a Typhoon FLA 9500 (GE Healthcare), and quantified using ImageQuant 5.2 (GE Healthcare).

### Binding analysis

Target binding kinetics were assessed by adding a trace amount of radiolabeled target RNA to varying concentrations (10-25 nM) of the assembled dCas13d-crRNA complex. These reactions were performed in a buffer containing 50 mM Tris-HCl, 150 mM NaCl, 1 mM DTT, 20 mM EDTA, and 0.2 mg/ml molecular-grade BSA, similar to the cleavage reaction conditions. At various time points, 2 µL aliquots were taken and added to 4 µL of an ice-cold chase solution (reaction buffer with 100 nM unlabeled target RNA, 15% glycerol, and xylene cyanol) and placed on ice. Control reactions, where chase target RNA and labeled target RNA were premixed and then added to the Cas13d complex, confirmed that the chase solution effectively competed against the labeled target RNA. After adding the chase solution, aliquots were kept on ice and then loaded onto a 15% native gel run at 4 °C. The gels were exposed overnight and analyzed as previously described. The association rate constant (k_on_) was determined from the slope of the observed rate constant versus Cas13d concentration. In the binding reactions for mutant Cas13d proteins, measurements were performed under Mg^2+^-free conditions without preparing a dead variant, and the binding rates were normalized to the without Mg^2+^ binding rate of the wild-type Cas13d.

### Cryo-EM sample preparation and data acquisition

Purified Cas13d complexes were diluted to 1 µM in 25 mM Tris pH 7.5, 200 mM NaCl, 5% glycerol, and 1 mM DTT. Samples were deposited on an Ultra Au foil R 1.2//1.3 grid (Quantifoil) that was plasma-cleaned for 1.5 min (Gatan Solarus 950). Excess liquid was blotted away for 4 s in a Vitrobot Mark IV (FEI) operating at 4 °C and 100% humidity before being plunge-frozen into liquid ethane. Data were collected on a Krios cryo-transmission electron microscope (TEM; Thermo Fisher Scientific) operating at 300 kV, equipped with a Gatan Biocontinuum Imaging Filter and a K3 direct electron detector camera (Gatan). Movies were collected using SerialEM at a pixel size of 0.8332 Å with a total exposure dose of 80 *e*^−^*/*Å^2^ and a defocus range of −1.2 to −2.2 µm[56]. To address a preferred orientation noticed during the collection of the C4A mismatch sample, subsequent datasets were gathered at a 30°tilt.

### Data analysis and model building

Real-time CTF correction, motion correction, and particle picking were executed using cryoSPARC Live[57]. Further data processing, including 2D classification, 3D ab initio, heterogeneous refinements, 3D variation analysis, 3D classification, and homogeneous refinements, occurred with cryoSPARC. A full description of the cryo-EM data processing workflows can be found in Figure S3-S6. A published EsdCas13d structure (PDB: 6E9F [https://doi.org/10.2210/pdb6E9F/pdb]) was docked into cryo-EM density maps using Chimera before being refined in Coot, ISOLDE, and PHENIX[18, 58–60]. Full cryo-EM data collection and refinement statistics can be found in Tables 3-6.

## Data Availability

Cryo-EM density maps have been deposited in the Electron Microscopy Data Bank (EMDB) under accession codes EMDB-47892 (binary complex), EMDB-47902 (active state), EMDB-47903 (intermediate state), EMDB-47904 (C4A intermediate state), EMDB-47905 (U10G intermediate state), EMDB-47906 (U10G inactive state), EMDB-47907 (U10G active state), and EMDB-47908 (C4A inactive state). Atomic coordinates have been deposited in the Protein Data Bank (PDB) under accession codes 9EBU (binary complex), 9EC8 (active state), 9EC9 (intermediate state), 9ECA (C4A intermediate state), 9ECB (U10G intermediate state), 9ECC (U10G inactive state), 9ECD (U10G active state), and 9ECE (C4A inactive state). All other data supporting the findings of this study are available from the corresponding author upon reasonable request.

## Declarations

## Author Contributions

C.-W.C., S.S., H.-C.K., and I.J.F. conceived the project. C.-W.C., H.-C.K., and S.S. performed protein purification, complex assembly, and biochemical experiments. D.S. cloned and purified some proteins. C.-W.C. performed cryo-EM sample preparation, data collection, and structural analysis. S.S. performed all kinetic analyses. C.A. and R.R. assisted with experimental design and data analysis. C.-W.C., S.S., and I.J.F. prepared the figures and wrote the manuscript with input from all coauthors. I.J.F. secured the funding and supervised the project.

## Acknowledgments

We thank all members of the Finkelstein lab for helpful discussions. Cryo-EM data collection was performed at the University of Texas at Austin Cryo-EM Facility. This work was supported by the Welch Foundation grant F-1808 (to I.J.F.), a generous gift from Tito’s Handmade Vodka, and a Spark Catalyst grant from the College of Natural Sciences at the University of Texas at Austin.

