## Supplementary material for "Structural Basis for Target Discrimination and Activation by Cas13d": Supplental Information

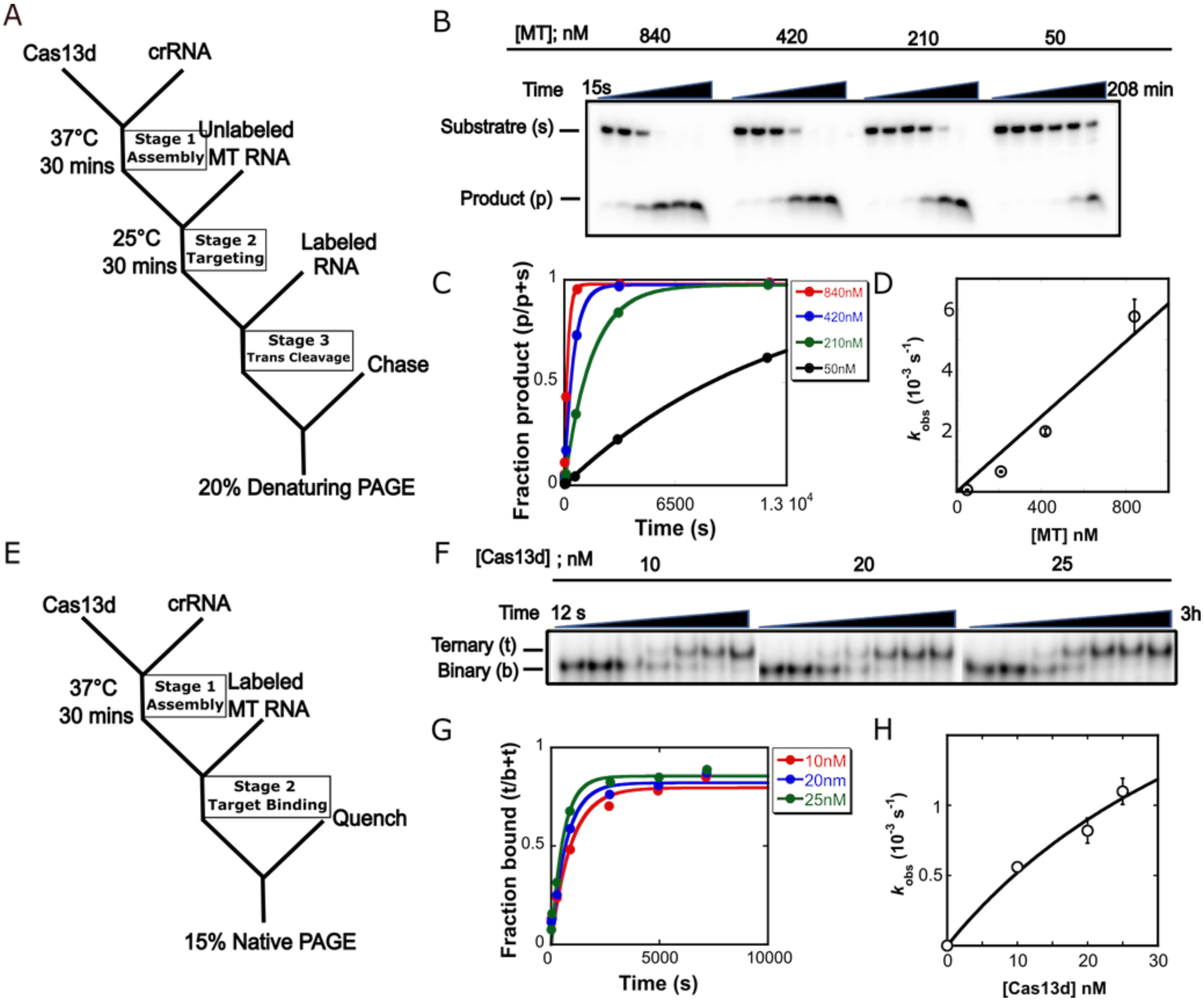

**Figure S1: Quantitative substrate cleavage and binding kinetics.** (A) Schematic of the *trans* cleavage assay with a 10 nt substrate. Stage 1: Cas13d assembles with crRNA. Stage 2: the assembled complex binds to a matched target. Stage 3: activated Cas13d cleaves the radiolabeled *trans* substrate. (B) Representative gel with varying concentrations of the activated Cas13d complex. The reaction initiates when the substrate RNA is added, and is quenched at various time points (15 s, 30 s, 105 s, 12 min, 55 min, 208 min). (C) Quantification of cleavage gels were fit to pseudo-first-order rate equations (solid lines) for the indicated activated RNP concentrations. (D) Second-order rate and maximal first-order rate constants from a hyperbolic fit to the observed rate constants from (C). (E) Schematic of the binding assay for Cas13d using a matched target RNA (40 nt). Stage 1: Cas13d assembles with crRNA. Stage 2: the assembled complex binds to the fully matched target. (F) Representative gel showing time courses for Cas13d binding to trace radiolabeled MT RNA. Time points: 12 s, 90 s, 9 min, 29 min, 1.5 h, 3 h. (G) Time-course data from the gel with fits to pseudo-first-order rate equations. (H) Second-order and maximal first-order rate constants from a hyperbolic fit to the data from (G). Results are presented as the mean with SEM from three independent experiments.

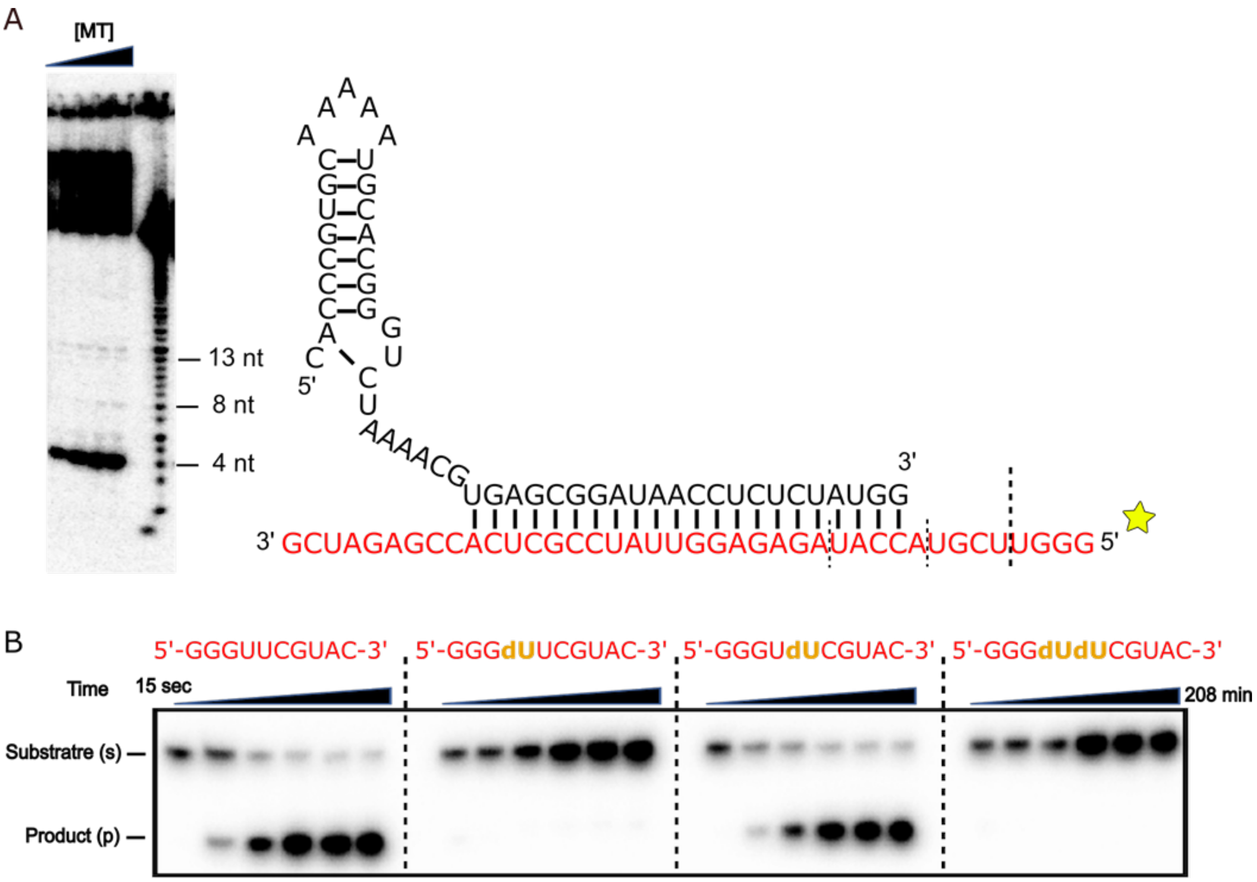

**Figure S2: Cas13d cleaves between UU dinucleotides.** (A) Cleavage of a radiolabeled matched target (MT) RNA (red) by Cas13d activated with an unlabeled MT RNA. Ladder: alkaline-hydrolyzed MT RNA. The cleavage products are marked by dashed lines. (B) Cleavage of a 10 nt *trans* substrate by activated Cas13d. Full cleavage requires a 5'-U ribose nucleotide; a deoxyribose sugar in this position inhibits all cleavage. The reaction is initiated by the addition of substrate RNA and quenched at different time points (15 s, 8 min, 30 min, 1 h, 2.5 h, 3.4 h)

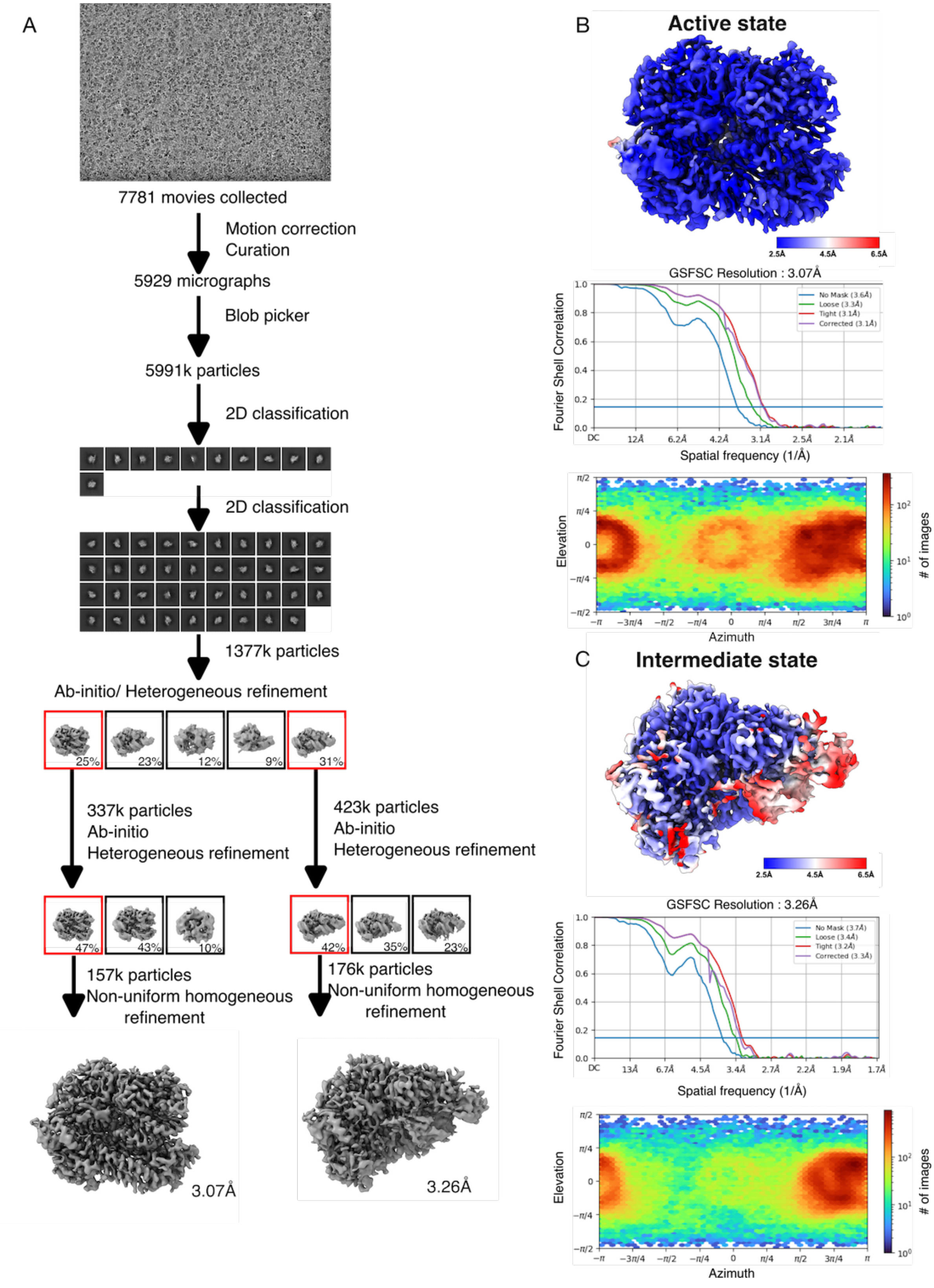
A
7781 movies collected
Motion correction
Curation
5929 micrographs
Blob picker
5991k particles
2D classification
2D classification
1377k particles
Ab-initio/ Heterogeneous refinement
25%
23%
12%
9%
31%
337k particles
Ab-initio
Heterogeneous refinement
423k particles
Ab-initio
Heterogeneous refinement
47%
43%
10%
42%
35%
23%
157k particles
Non-uniform homogeneous refinement
176k particles
Non-uniform homogeneous refinement
3.07Å
3.26Å
B
Active state
2.5Å
4.5Å
6.5Å
GSFSC Resolution : 3.07Å
Fourier Shell Correlation
Spatial frequency (1/Å)
No Mask (3.6Å)
Loose (3.3Å)
Tight (3.1Å)
Corrected (3.1Å)
Elevation
Azimuth
### of images
C
Intermediate state
2.5Å
4.5Å
6.5Å
GSFSC Resolution : 3.26Å
Fourier Shell Correlation
Spatial frequency (1/Å)
No Mask (3.7Å)
Loose (3.4Å)
Tight (3.2Å)
Corrected (3.3Å)
Elevation
Azimuth
### of images

**Figure S3: Cryo-EM data collection, analysis, and resolution estimation of active state and intermediate state of Cas13d-crRNA-matched-target RNA ternary complex** (A) Workflow of data collection and analysis of Cas13d-crRNA-matched-target RNA ternary complex[1, 2]. (B) Refined map of the active state of Cas13d-crRNA-matched-target complex colored by local resolution (top). Fourier Shell Correlation (FSC) curves for cross-validation between two half maps of the active state of Cas13d-crRNA-matched-target RNA ternary complex. Resolutions were estimated at FSC=0.143 (middle). Euler diagrams showing orientation distributions of the particles used in the final 3D reconstruction (bottom). (C) Refined map of the intermediate state of Cas13d-crRNA-matched-target complex colored by local resolution (top). Fourier Shell Correlation (FSC) curves for cross-validation between two half maps of the active state of Cas13d-crRNA-matched-target RNA ternary complex. Resolutions were estimated at FSC=0.143 (middle). Euler diagrams showing orientation distributions of the particles used in the final 3D reconstruction (bottom).

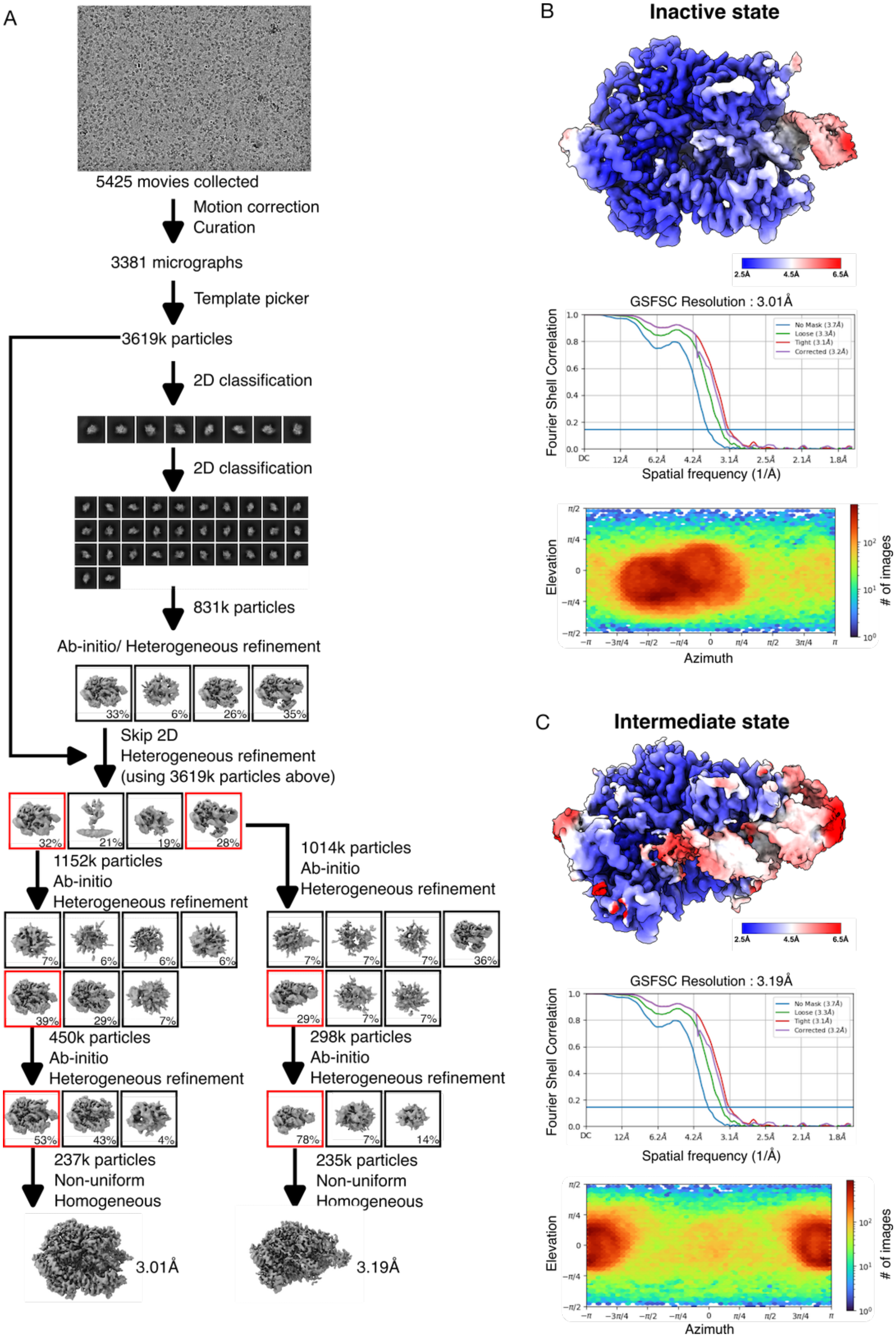
A
5425 movies collected
Motion correction
Curation
3381 micrographs
Template picker
3619k particles
2D classification
2D classification
831k particles
Ab-initio/ Heterogeneous refinement
33%
6%
26%
35%
Skip 2D
Heterogeneous refinement
(using 3619k particles above)
32%
21%
19%
28%
1152k particles
Ab-initio
Heterogeneous refinement
1014k particles
Ab-initio
Heterogeneous refinement
450k particles
Ab-initio
Heterogeneous refinement
298k particles
Ab-initio
Heterogeneous refinement
237k particles
Non-uniform
Homogeneous
235k particles
Non-uniform
Homogeneous
3.01Å
3.19Å
B
Inactive state
2.5Å
4.5Å
6.5Å
GSFSC Resolution : 3.01Å
Fourier Shell Correlation
Spatial frequency (1/Å)
Elevation
Azimuth
### of images
C
Intermediate state
GSFSC Resolution : 3.19Å
Fourier Shell Correlation
Spatial frequency (1/Å)
Elevation
Azimuth
### of images

**Figure S4: Cryo-EM data collection, analysis, and resolution estimation of the active and intermediate Cas13d ternary complexes.** (A) Workflow of data collection and analysis of Cas13d-crRNA-matched-target RNA ternary complex[1, 2].(B) Refined map of the active state of Cas13d-crRNA-matched-target complex colored by local resolution (top). Fourier Shell Correlation (FSC) curves for cross-validation between two half maps of the active state of Cas13d-crRNA-matched-target RNA ternary complex. Resolutions were estimated at FSC=0.143 (middle). Euler diagrams showing orientation distributions of the particles used in the final 3D reconstruction (bottom).(C) Refined map of the intermediate state of Cas13d-crRNA-matched-target complex colored by local resolution (top). Fourier Shell Correlation (FSC) curves for cross-validation between two half maps of the active state of Cas13d-crRNA-matched-target RNA ternary complex. Resolutions were estimated at FSC=0.143 (middle). Euler diagrams showing orientation distributions of the particles used in the final 3D reconstruction (bottom).

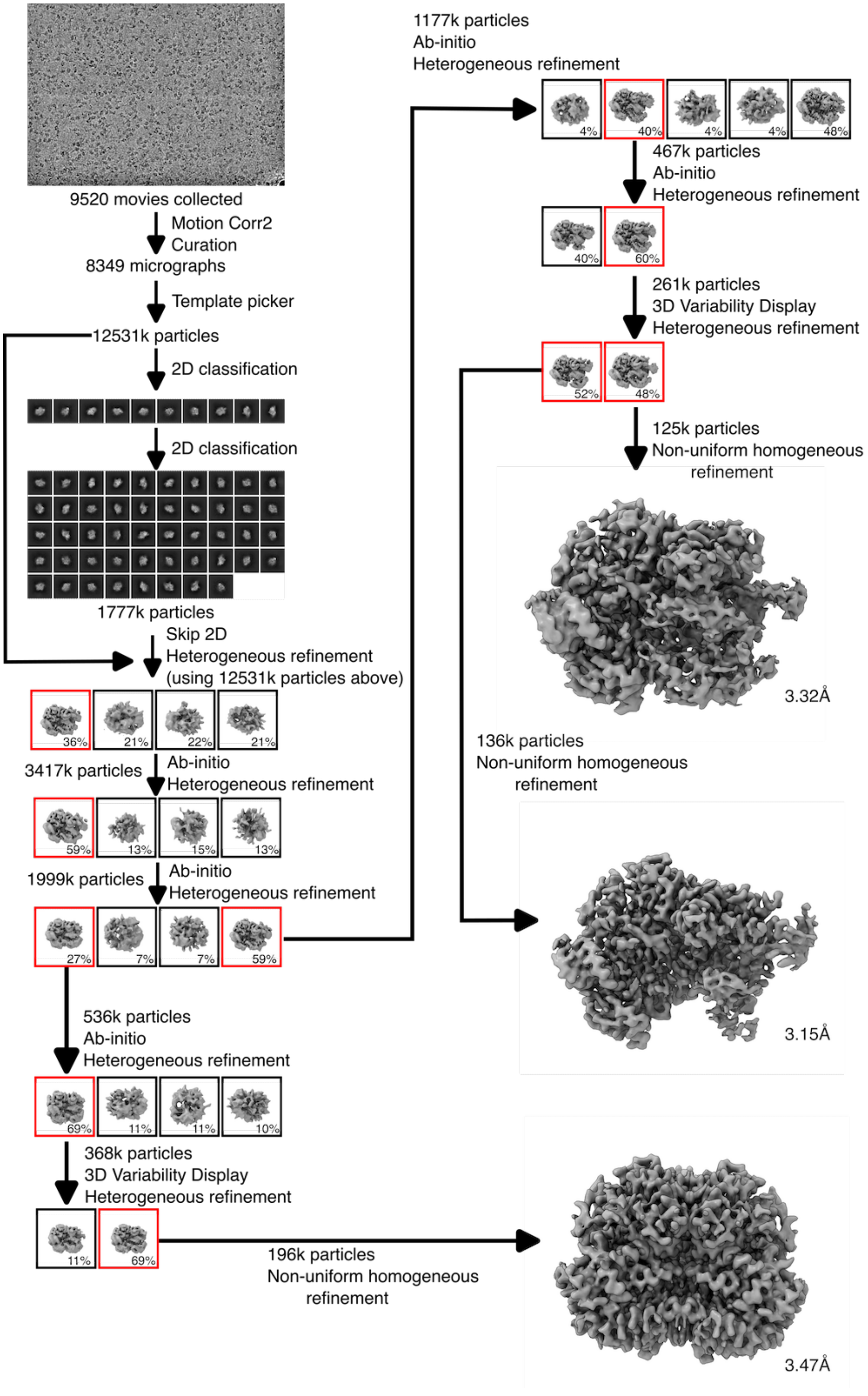

**Figure S5: Cryo-EM data collection and analysis of Cas13d-crRNA-U10G-target RNA ternary complex.** Workflow of data collection and analysis of Cas13d-crRNA-U10G-target RNA ternary complex[1, 2].

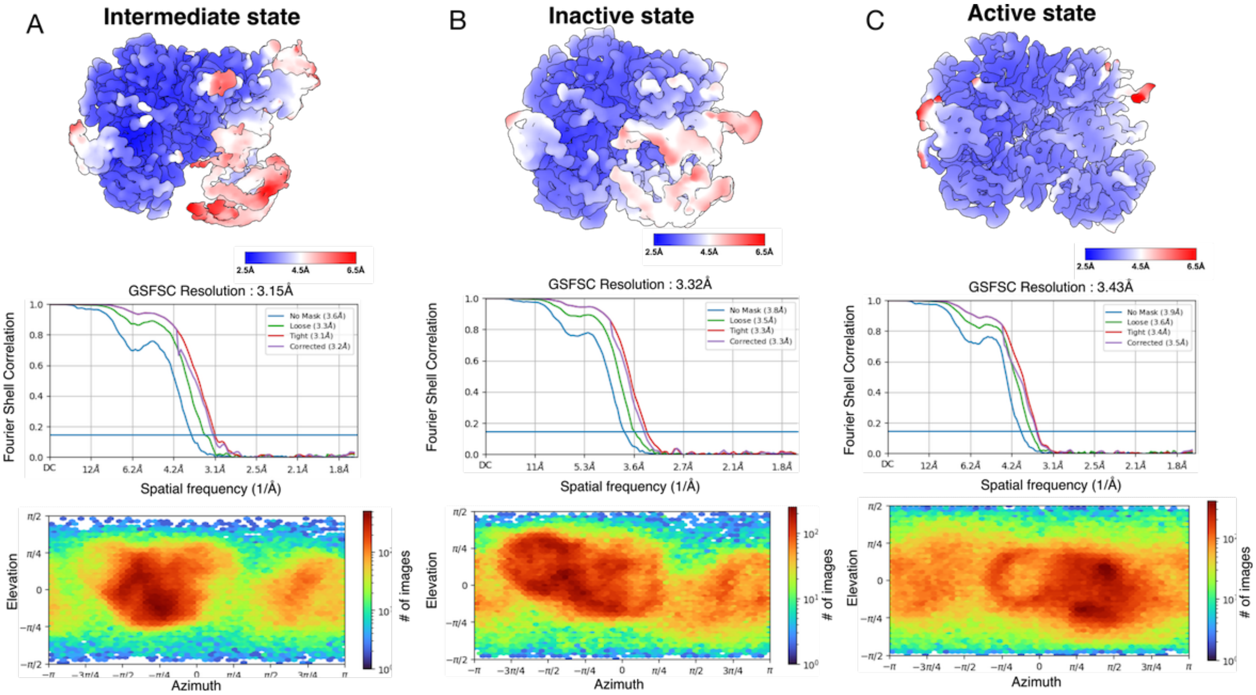

**Figure S6: Cryo-EM data analysis of Cas13d-crRNA-U10G-target RNA ternary complex.** Refined map of the (A) intermediate state, (B) inactive state, (C) active state of Cas13d-crRNA-U10G-target complex colored by local resolution (top). Fourier Shell Correlation (FSC) curves for cross-validation between two half maps of the active state of Cas13d-crRNA-matched-target RNA ternary complex. Resolutions were estimated at FSC=0.143 (middle). Euler diagrams showing orientation distributions of the particles used in the final 3D reconstruction (bottom).

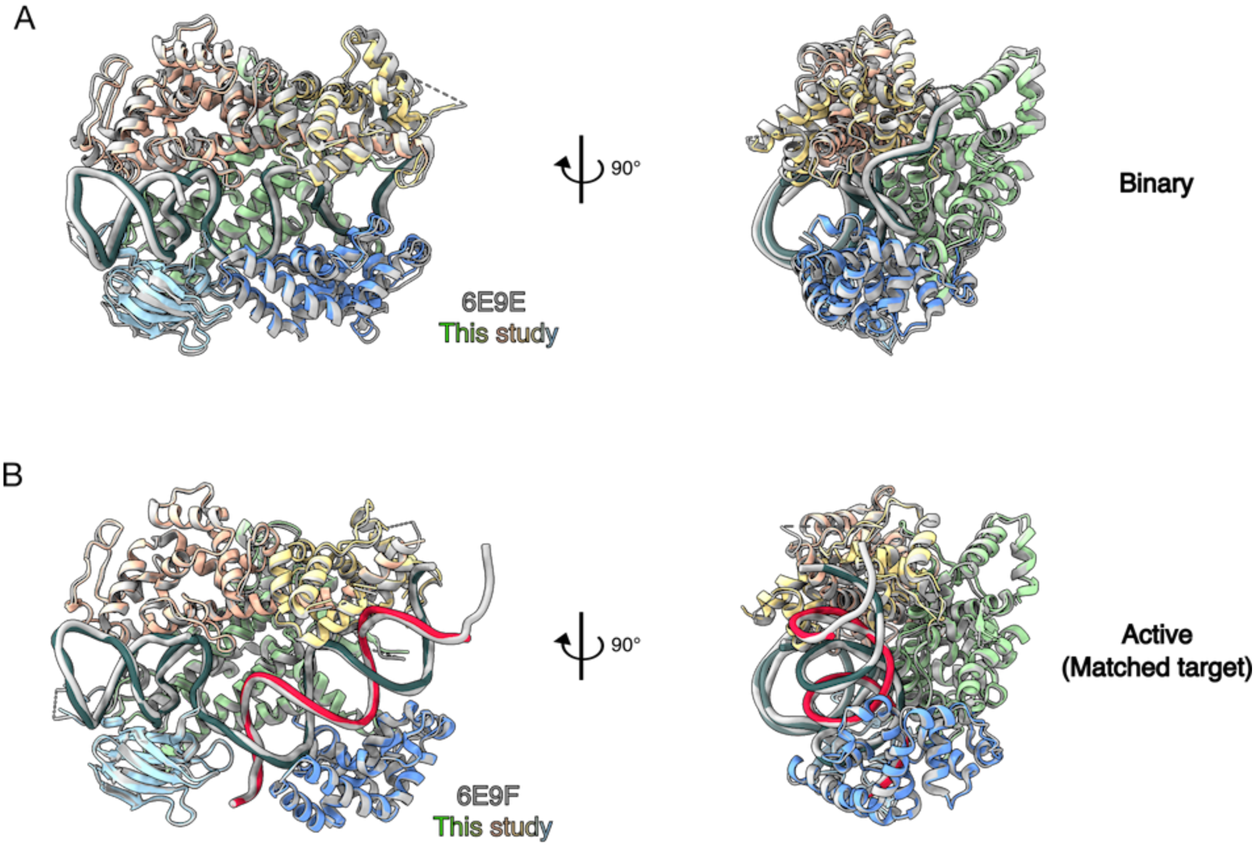

**Figure S7: Comparison to published EsCas13d structures.** (A) Superposition of binary EsCas13d structures. Gray: 6E9E from Zhang et al.[3]. Colored: this study. The $C\alpha$-$C\alpha$-RMSD is 1.327 Å. (B) Superposition of active EsCas13d structures. Gray: 6E9F from Zhang et al.[3]. Colored: this study. The $C\alpha$-$C\alpha$-RMSD is 0.700 Å.

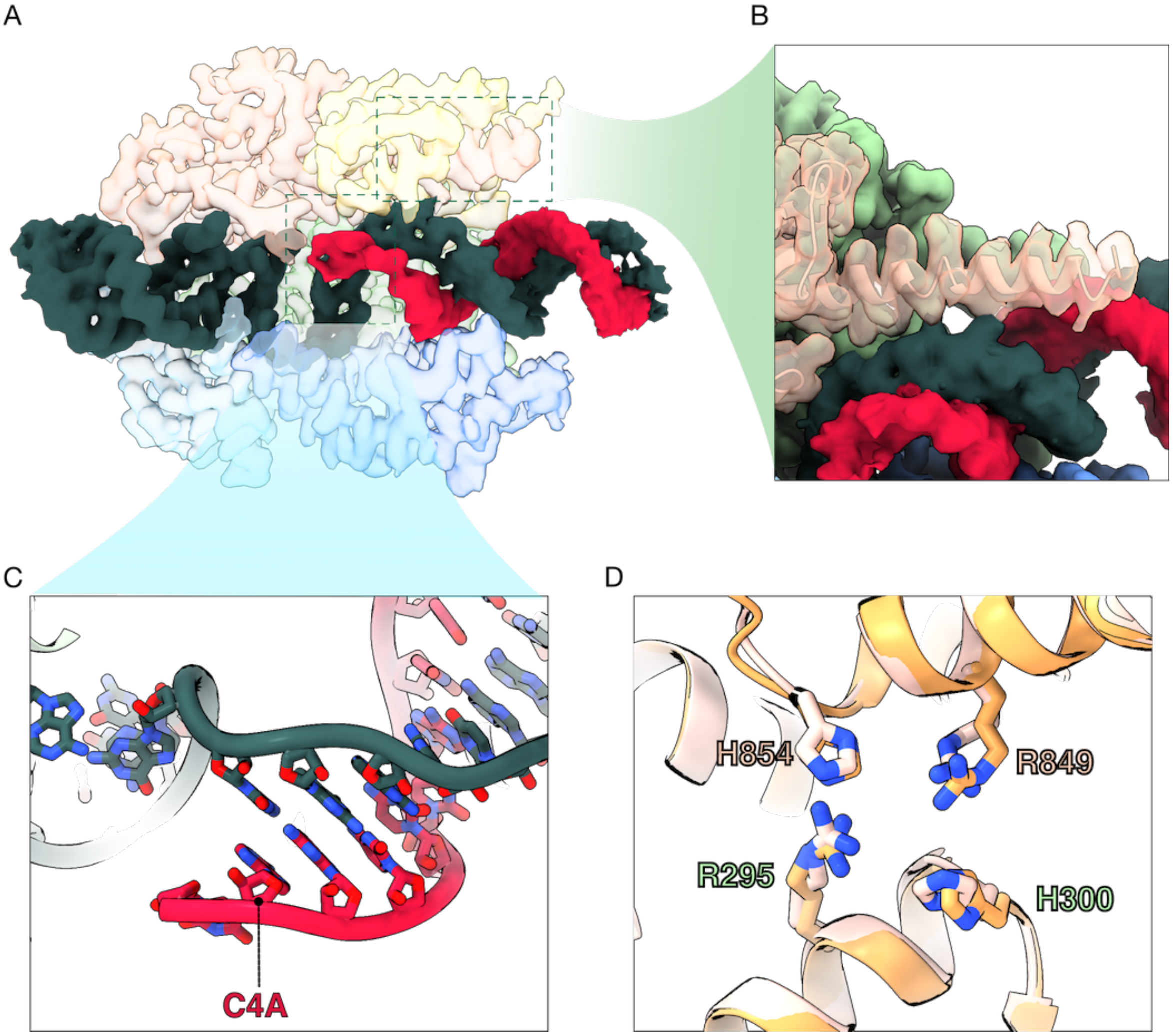

**Figure S8: Structural features of the inactive state.** (A) EM density of the Cas13d ternary inactive state (C4A target), colored by domains. Green: crRNA, Red: partial target RNA, Pink: HEPN2 domain, Yellow: Helical-2 domain, Light blue: NTD domain, Dark blue: Helical-1 domain. (B) Enlarged view of an elongated $\alpha$-helix in the HEPN2 domain. (C) Enlarged view of the RNA duplex with missing base pairing at the proximal end, near the C4A mismatch. (D) Enlarged view of the nuclease active site in the inactive state (pink) overlapped with the binary structure from this study (orange).

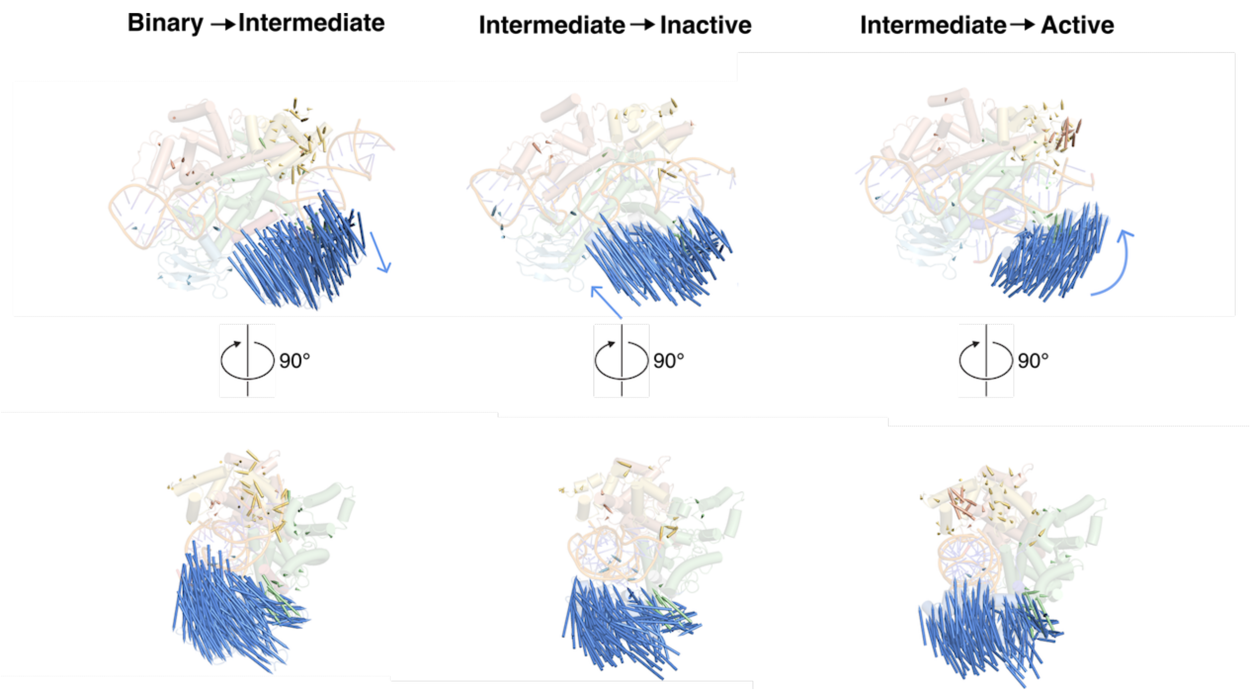

**Figure S9: Conformational changes between the binary, intermediate, inactive, and active states.** Transparent cartoon representations of binary (left), intermediate (middle), and active (right) states of Cas13d with vector arrows indicating the conformational changes of Cas13d upon target binding and activation. Blue arrows represent the vector direction of the Helical-1 domain.

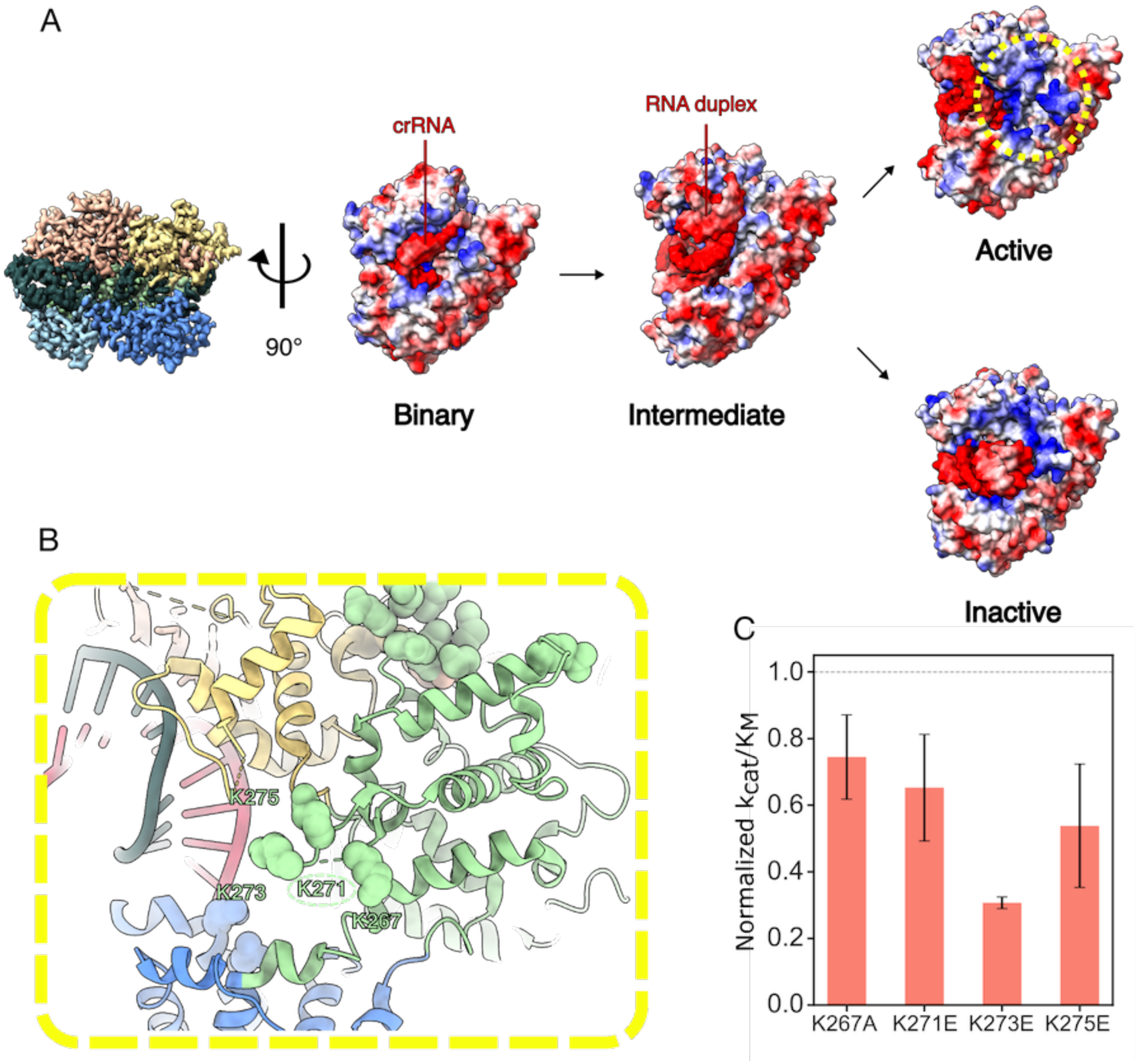

**Figure S10: Cas13d activation reveals a positive-charge tunnel between Helical-2 and HEPN1 domain.** (A) Electrostatic surfaces of Cas13d from different states. The dashed circle highlights the exposed positive charge in the active state. (B) Enlarged view of the exposed HEPN1 domain positive residues in the active state. (C) Cleavage activity of the indicated mutants in the positive charge tunnel. Error bars: mean and SEM from 3 independent experiments.

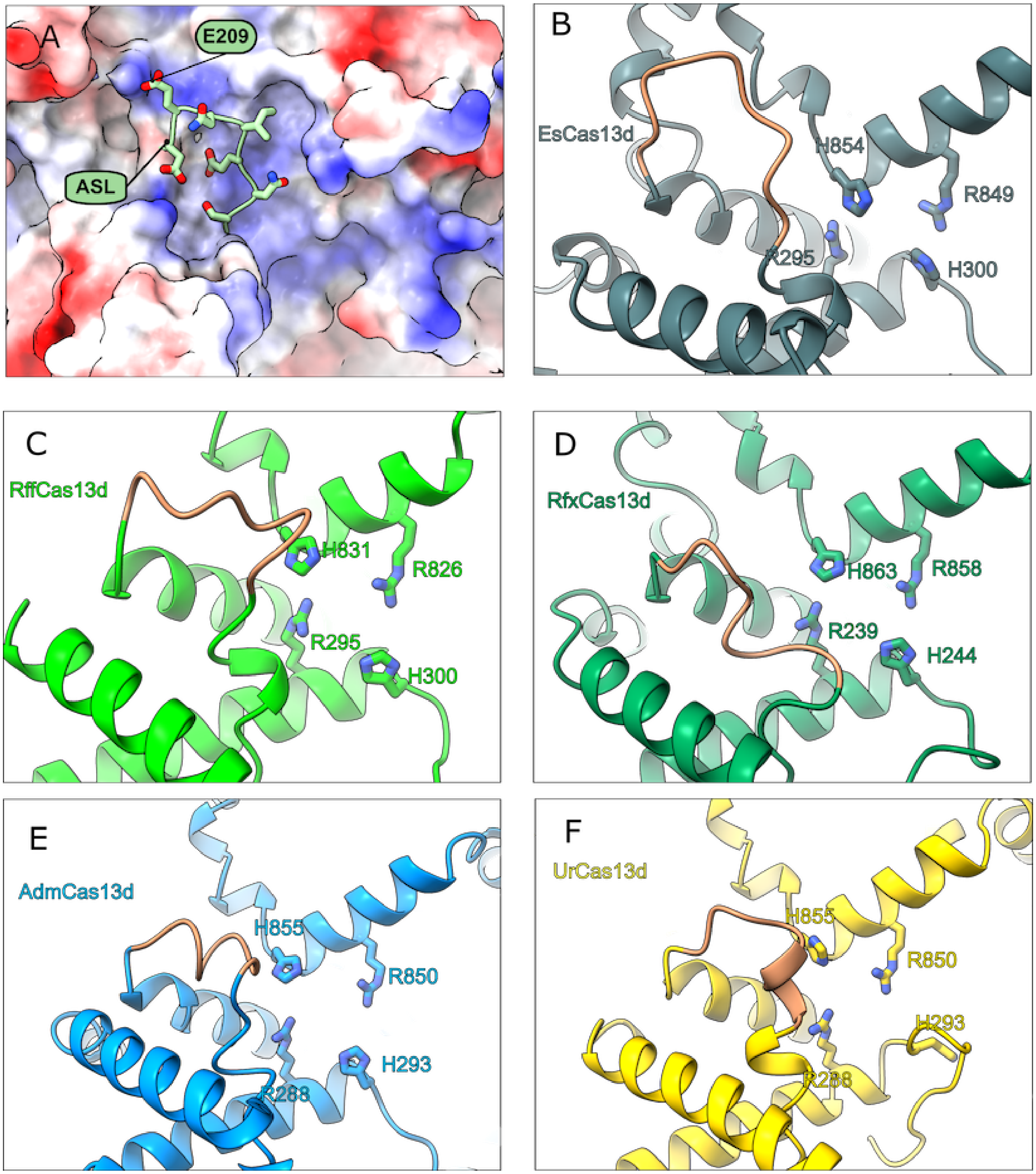

**Figure S11: AlphaFold3-predicted structures of multiple Cas13d orthologs suggest that the active site loop (ASL) is conserved.** (A) Enlarged views of ASL adjacent to the active site are shown on the electrostatic surface. (B-F) ASL and active sites in the Cas13 orthologs. Salmon: ASLs. The key active site residues are indicated explicitly.

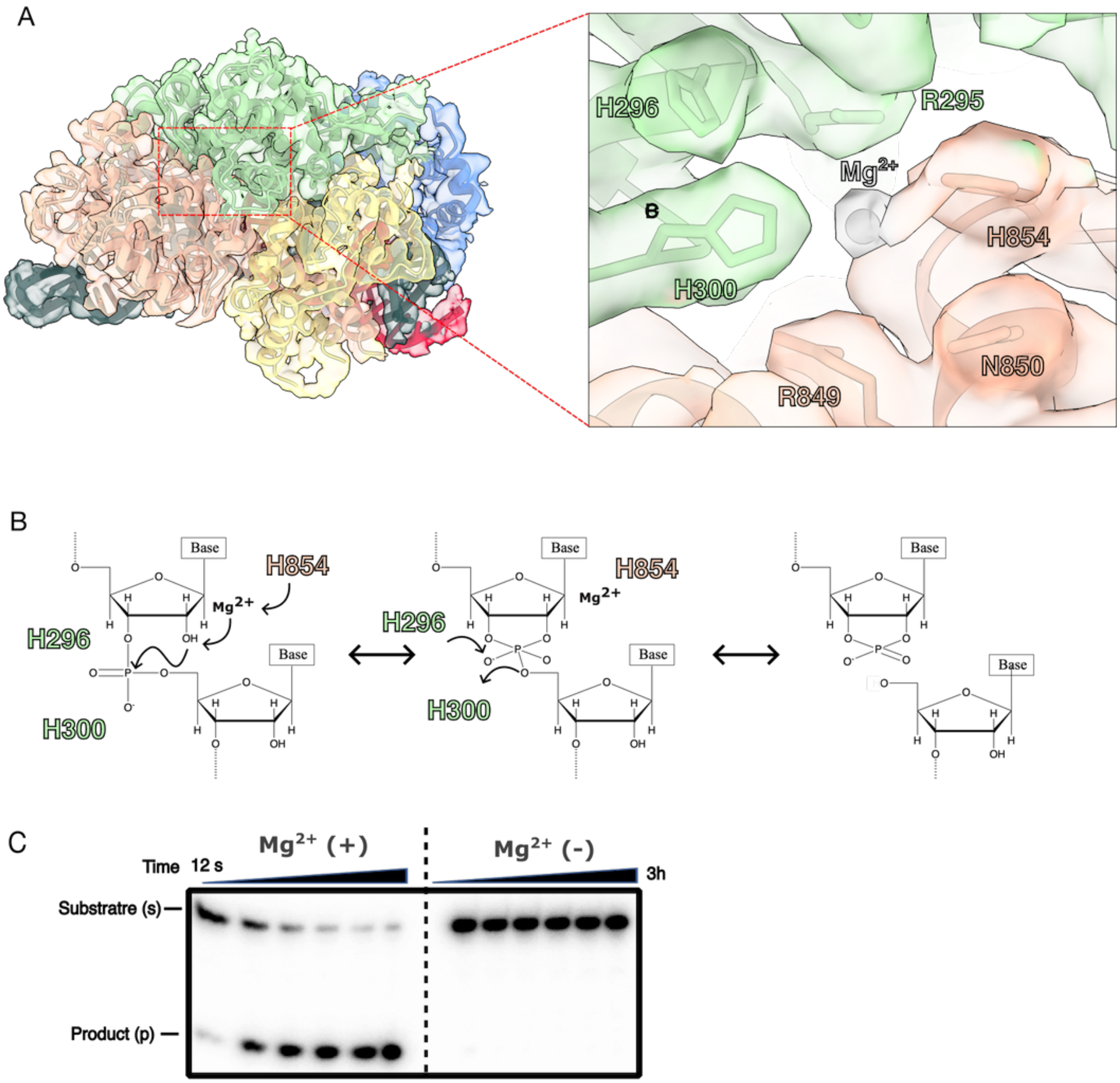

**Figure S12: A potential $Mg^{2+}$ ion is involved in cleavage mechanism.** (A) Cryo-EM structure of the active site highlighting the $Mg^{2+}$ ion (grey). A potential $Mg^{2+}$ ion is coordinated by the HEPN nuclease motif (R$\phi$XXXH). (B) Proposed mechanism of trans RNA cleavage by acid-base catalysis. H854 acts as the general base, abstracting a proton from the 2'-OH adjacent to the scissile phosphate, with $Mg^{2+}$ facilitating this process. H300 or H296 then serves as a general acid, protonating the 5'-oxygen leaving group. (C) Gel showing time-course of trans-RNA cleavage in the presence (left) and absence (right) of $Mg^{2+}$. Cleavage occurs more rapidly with $Mg^{2+}$, while little to no cleavage is observed without $Mg^{2+}$, indicating the essential role of $Mg^{2+}$ in the cleavage reaction.

#### Supplemental Tables

| Plasmid | Description | Source |
| --- | --- | --- |
| pIF1023 | 6xHis-TwinStrep-SUMO-Cas13d | Kuo et al. |
| pIF1024 | 6xHis-TwinStrep-SUMO-dCas13d | Kuo et al. |

**Table 1:** Plasmids used in this study[4].

| Sample | Sequence |
|---|---|
| R1_crRNA | CACCCGUGCAAAAAUGCAGGGGUCUAAAACUGAGCGGAUAACCUCUCUAUGG |
| R2_MT_kinetic | GGGUUCGUACCAUAGAGAGGUUAUCCGCUCACCGAGAUCG |
| R3_C4A_kinetic | GGGUUCGUACCAUAGAGAGGUUAUCCGAUCACCGAGAUCG |
| R4_U10G_kinetic | GGGUUCGUACCAUAGAGAGGUGAUCCGCUCACCGAGAUCG |
| R5_MT_structure | CGUACCAUAGAGAGGUUAUCCGCUCACCGA |
| R6_C4A_structure | CGUACCAUAGAGAGGUUAUCCGAUCACCGA |
| R7_U10G_structure | CGUACCAUAGAGAGGUGAUCCGCUCACCGA |
| R8_Target-1(T1) | GGGUUCGUACCAUAGAGAGGUUAUCCGCUC |
| R9_Target-2(T2) | GGGUUCGUACCAUAGAGAGGUUAUCCGCUCA |
| R10_Target-3(T3) | GGGUUCGUACCAUAGAGAGGUUAUCCGCUCAC |
| R11_Target-4(T4) | GGGUUCGUACCAUAGAGAGGUUAUCCGCUCACC |
| R12_Target-5(T5) | GGGUUCGUACCAUAGAGAGGUUAUCCGCUCACCGA |
| R13_trans_substrate | GGGUUCGUAC |
| R14_trans_4'dU_substrate | GGGdUUCGUAC |
| R15_trans_5'dU_substrate | GGGUdUCGUAC |
| R16_trans_4',5'dU_substrate | GGGdUdUCGUAC |

**Table 2:** RNA sequences used in this study.

| | Binary complex of EsCas13d (EMDB-xxxx) (PDB-9EBU) |
|---|---|
| **Data collection and Processing** | |
| Magnification | 29,000 |
| Voltage (kV) | 200 |
| Electron exposure ($e^{-}$/$Å^2$) | 80 |
| Defocus range ($\mu$m) | -1.2 to -2.2 |
| Pixel size (Å) | 0.81 |
| Symmetry imposed | C1 |
| Initial particle images (no.) | 1,248k |
| Final particle images (no.) | 192k |
| Map resolution (Å) | 3.06 |
| FSC threshold = 0.143 | |
| Map resolution range (Å) | 3-7 |
| **Refinement** | |
| Initial model used (PDB code) | 6E9E |
| Model resolution (Å) | 3.1 |
| FSC threshold | 0.5 |
| Model composition | |
| Non-hydrogen atoms | 8097 |
| Protein residues | 860 |
| Nucleotides | 49 |
| B factors ($Å^2$) | |
| Protein | 44.30 |
| Ligand | 58.66 |
| R.m.s. deviations | |
| Bond lengths (Å) | 0.005 |
| Bond angles (°) | 0.631 |
| Validation | |
| MolProbity score | 2.38 |
| Clashscore | 7.88 |
| Poor rotamers (%) | 3.73 |
| Ramachandran plot | |
| Favored (%) | 97.64 |
| Allowed (%) | 2.36 |
| Disallowed (%) | 0.00 |

**Table 3:** EsCas13d binary complex Cryo-EM data collection and processing statistics

| | EsCas13d with matched target | |
|---|---|---|
| | Intermediate state (EMDB-47903) (PDB-9EC9) | Active state (EMDB-47902) (PDB-9EC8) |
| **Data collection and Processing** | | |
| Magnification | 28,000 | 28,000 |
| Voltage (kV) | 300 | 300 |
| Electron exposure ($e^{-}/Å^2$) | 80 | 80 |
| Defocus range ($\mu$m) | -1.2 to -2.2 | -1.2 to -2.2 |
| Pixel size (Å) | 0.8332 | 0.8332 |
| Symmetry imposed | C1 | C1 |
| Initial particle images (no.) | 1377k | 1377k |
| Final particle images (no.) | 157k | 176k |
| Map resolution (Å) | 3.07 | 3.26 |
| FSC threshold | 0.143 | 0.143 |
| Map resolution range (Å) | 3 - 7 | 3 - 7 |
| **Refinement** | | |
| Initial model used (PDB code) | 6E9F | 6E9F |
| Model resolution (Å) | 3.3 | 3.1 |
| FSC threshold | 0.5 | 0.5 |
| Model composition | | |
| Non-hydrogen atoms | 7006 | 8663 |
| Protein residues | 676 | 861 |
| Nucleotides | 69 | 75 |
| B factors ($Å^2$) | | |
| Protein | 72.85 | 90.90 |
| Nucleotide | 130.50 | 103.56 |
| R.m.s. deviations | | |
| Bond lengths (Å) | 0.004 | 0.003 |
| Bond angles (°) | 0.864 | 0.567 |
| Validation | | |
| MolProbity score | 1.54 | 1.38 |
| Clashscore | 7.39 | 6.95 |
| Poor rotamers (%) | 0.33 | 0.26 |
| Ramachandran plot | | |
| Favored (%) | 97.58 | 98.12 |
| Allowed (%) | 2.42 | 1.88 |
| Disallowed (%) | 0.00 | 0.00 |

**Table 4:** EsCas13d-MT ternary complex Cryo-EM data collection and processing statistics

| | EsCas13d with C4A target | |
|---|---|---|
| | Intermediate state (EMDB-47904) (PDB-9ECA) | Inactive state (EMDB-47908) (PDB-9ECE) |
| **Data collection and Processing** | | |
| Magnification | 28,000 | 28,000 |
| Voltage (kV) | 300 | 300 |
| Electron exposure ($e^{-}/Å^2$) | 80 | 80 |
| Defocus range ($\mu$m) | -1.2 to -2.2 | -1.2 to -2.2 |
| Pixel size (Å) | 0.8332 | 0.8332 |
| Symmetry imposed | C1 | C1 |
| Initial particle images (no.) | | |
| Final particle images (no.) | | |
| Map resolution (Å) | | |
| FSC threshold | | |
| Map resolution range (Å) | | |
| **Refinement** | | |
| Initial model used (PDB code) | | |
| Model resolution (Å) | 3.4 | 3.1 |
| FSC threshold | 0.5 | 0.5 |
| Model composition | | |
| Non-hydrogen atoms | 6918 | 8299 |
| Protein residues | 670 | 835 |
| Nucleotides | 67 | 68 |
| B factors ($Å^2$) | | |
| Protein | 81.25 | 91.82 |
| Nucleotide | 146.62 | 144.89 |
| R.m.s. deviations | | |
| Bond lengths (Å) | 0.003 | 0.003 |
| Bond angles (°) | 0.769 | 0.550 |
| Validation | | |
| MolProbity score | 1.92 | 1.67 |
| Clashscore | 9.42 | 9.77 |
| Poor rotamers (%) | 1.67 | 1.60 |
| Ramachandran plot | | |
| Favored (%) | 96.33 | 98.05 |
| Allowed (%) | 3.67 | 1.95 |
| Disallowed (%) | 0.00 | 0.00 |

**Table 5:** EsCas13d-C4A ternary complex cryo-EM data collection and processing statistics

| | | EsCas13d with U10G | |
|---|---|---|---|
| | Intermediate (EMDB-47905) (PDB-9ECB) | Inactive state (EMDB-47906) (PDB-9ECC) | Active state (EMDB-47907) (PDB-9ECD) |
| **Data collection and Processing** | | | |
| Magnification | | | |
| Voltage (kV) | | | |
| Electron exposure ($e^{-}/Å^2$) | | | |
| Defocus range ($\mu$m) | | | |
| Pixel size (Å) | | | |
| Symmetry imposed | | | |
| Initial particle images (no.) | | | |
| Final particle images (no.) | | | |
| Map resolution (Å) | | | |
| FSC threshold | | | |
| Map resolution range (Å) | | | |
| **Refinement** | | | |
| Initial model used (PDB code) | | | |
| Model resolution (Å) | 3.1 | 3.1 | 3.6 |
| FSC threshold | 0.5 | 0.5 | 0.5 |
| Model composition | | | |
| Non-hydrogen atoms | 8162 | 8120 | 8143 |
| Protein residues | 832 | 831 | 821 |
| Nucleotides | 62 | 61 | 66 |
| B factors ($Å^2$) | | | |
| Protein | 70.79 | 88.13 | 118.92 |
| Nucleotide | 102.68 | 142.31 | 140.24 |
| R.m.s. deviations | | | |
| Bond lengths (Å) | 0.003 | 0.003 | 0.003 |
| Bond angles (°) | 0.823 | 0.616 | 0.593 |
| Validation | | | |
| MolProbity score | 1.49 | 1.60 | 1.60 |
| Clashscore | 6.01 | 7.65 | 6.56 |
| Poor rotamers (%) | 1.61 | 1.74 | 2.03 |
| Ramachandran plot | | | |
| Favored (%) | 97.07 | 98.29 | 98.02 |
| Allowed (%) | 2.80 | 1.71 | |
| 1.98 Disallowed (%) | 0.12 | 0.00 | 0.00 |

**Table 6:** EsCas13d-U10G ternary complex cryo-EM data collection and processing statistics.
